# Infection and sensitization reveal stimulus-specific immune remodeling in aged skin

**DOI:** 10.64898/2026.09.03.749212

**Authors:** Morgan M. Severn, Anita Y. Voigt, Ashok Kumar Dhinakaran, Ruoyu Yang, Elizabeth Aiken, Soo-Yeon Kang, Sasan Jalili, Julia Oh

**Affiliations:** Department of Dermatology, Duke University School of Medicine, Durham, North Carolina, USA; The Jackson Laboratory for Genomic Medicine, Farmington, Connecticut, USA; Department of Immunology, School of Medicine, UConn Health, Farmington, CT, USA

**Author notes:** contributed equally (Morgan M. Severn, Anita Y. Voigt, Ashok Kumar Dhinakaran). **Corresponding author:** Julia Oh, Ph.D., Department of Dermatology Duke University School of Medicine, Durham, NC 27710 USA.

**Keywords:** skin aging, *Staphylococcus aureus* infection, tissue-resident immunity, microneedle patch sampling, immunosenescence, T cell response, barrier immunity

## Abstract

Aging is associated with progressive declines in skin barrier integrity and immune protection, contributing to increased susceptibility to bacterial and viral skin infections in older adults. However, how aged skin senses and responds to infection or barrier disruption remains poorly defined. Here, we characterized the age-associated cutaneous immune response to epicutaneous *Staphylococcus (S.) aureus* infection and ovalbumin-induced sensitization in mouse models. Bulk RNA-seq of infected tissue showed that transcriptional variation was primarily driven by infection, not age, suggesting that aged skin retains a broadly inducible response to microbial challenge. In contrast, microneedle patch (MNP) sampling of skin interstitial fluid, which provides cellular resolution, revealed age-dependent differences in local immune dynamics after *S. aureus* infection, including altered magnitude and kinetics of cellular recruitment, such as an attenuated cutaneous T cell response in older mice. MNP profiling further showed that ovalbumin sensitization elicited a localized immune program distinct from the *S. aureus* response, which was incompletely reflected in systemic measurements. Together, these data demonstrate that aging does not uniformly impair cutaneous immunity but instead is a context-dependent remodeling of local tissue immune dynamics.

**One-line summary:** Aging does not uniformly attenuate cutaneous immunity but instead remodels tissue-resident immune dynamics in a context-dependent manner depending on the skin challenge.

## INTRODUCTION

Aging is a complex systemic process shaped by intrinsic factors, lifestyle, and environmental exposures. The rapid expansion of the global older adult population has intensified the need to understand the mechanisms that support healthy aging and preserve tissue resilience. This is particularly important for barrier tissues such as the skin, which undergo progressive structural and immunologic changes that can increase susceptibility to injury, inflammation, infection, and numerous skin diseases.

Aged skin is exposed to intrinsic (i.e. reactive oxygen species) and extrinsic (i.e. ultraviolet radiation) factors that contribute to a progressive decline in barrier function and integrity^1,2^. Diminishing numbers of epidermal and dermal cells compromise the structural integrity of skin and increase susceptibility to injury and infection. Aging skin also undergoes a variety of innate and adaptive immunological changes which can impair or delay the magnitude, timing, and quality of responses to skin challenges^3,4^. Aging skin is additionally enriched in senescent keratinocytes and fibroblasts with senescence-associated secretory phenotypes, contributing to a proinflammatory cytokine milieu^4^. Together, these changes suggest that aged skin may not simply be immunologically diminished but may instead differentially respond to skin infection or barrier disruption.

Skin disorders are highly prevalent in older adults and range from benign to life threatening infections^3,5^. Bacterial and viral skin and soft tissue infections (SSTIs) are common in older adults, and are often attributed to changes in skin architecture, declining skin barrier properties over time, and impaired immunity^3,5,6^. *Staphylococcus (S.) aureus* is the leading cause of SSTIs in the United States (up to 74%)^7,8^, and circulating strains of multidrug resistant and methicillin resistant *S. aureus* (MRSA) are a major public health concern in community and hospital settings^7^. Older adults may be particularly vulnerable from this increased healthcare exposure, particularly in skilled nursing facilites^6,9,10^. Delayed wound healing and age-associated changes in cutaneous immunity likely further increase risk. However, few studies have mechanistically examined how aged skin senses and responds to *S. aureus* topical infection at the tissue level. Whether aged skin mounts diminished, delayed, or distinct responses to topical infection, and whether these are global or context-specific responses, remains unclear.

Here, we adapted a mouse model of epicutaneous infection^11,12^ to characterize age-associated changes in the cutaneous response to *S. aureus*. Because traditional studies primarily rely on endpoint analyses to assess local and systemic immune responses, they provide limited insight into the kinetics of local immune responses over time. To address this limitation, we implemented sampling microneedle patches (MNP) to longitudinally collect skin interstitial fluid and profile cytokine and immune cellular responses at the site of challenge and performed RNA-seq at endpoint to define broad infection resolution. This minimally invasive approach enables repeated assessment of tissue-level immune dynamics^13,14^ without requiring serial biopsy which would significantly alter the sampled area^14^, or animal sacrifice. We paired longitudinal MNP-based immune profiling with cross-sectional transcriptomic analysis of aged and young murine skin after *S. aureus* and ovalbumin-induced sensitization and defined longitudinal, age-dependent, tissue resident cytokine and cellular responses. Together, these approaches reveal that aging does not uniformly impair cutaneous immunity but instead remodels tissue immune dynamics in a context dependent manner.

## RESULTS

### Infection resolution is largely preserved across age, sex, barrier status, and genetic background

We first established an epicutaneous infection model of a well-characterized virulent *S. aureus* strain (USA300 LAC)^11,12^ to determine whether age, sex, genetic background and heterogeneity, or skin barrier status substantially altered the broad resolution of topical *S. aureus* infection (**Figure 1A**). Here, we modeled epicutaneous infection in four mouse strains covering a range of genetic diversity, including two inbred C57BL/6J and BALB/cJ strains and two outbred genetic diversity models “HET3” (CByB6F1/J x C3D2F1/J)/J and J:DO (diversity outbred), which is the progeny of eight crossbred founder strains^15^. *S. aureus* lesion severity was assessed by redness, scaliness, and skin integrity and tracked for one week post-infection.

**Figure 1:**
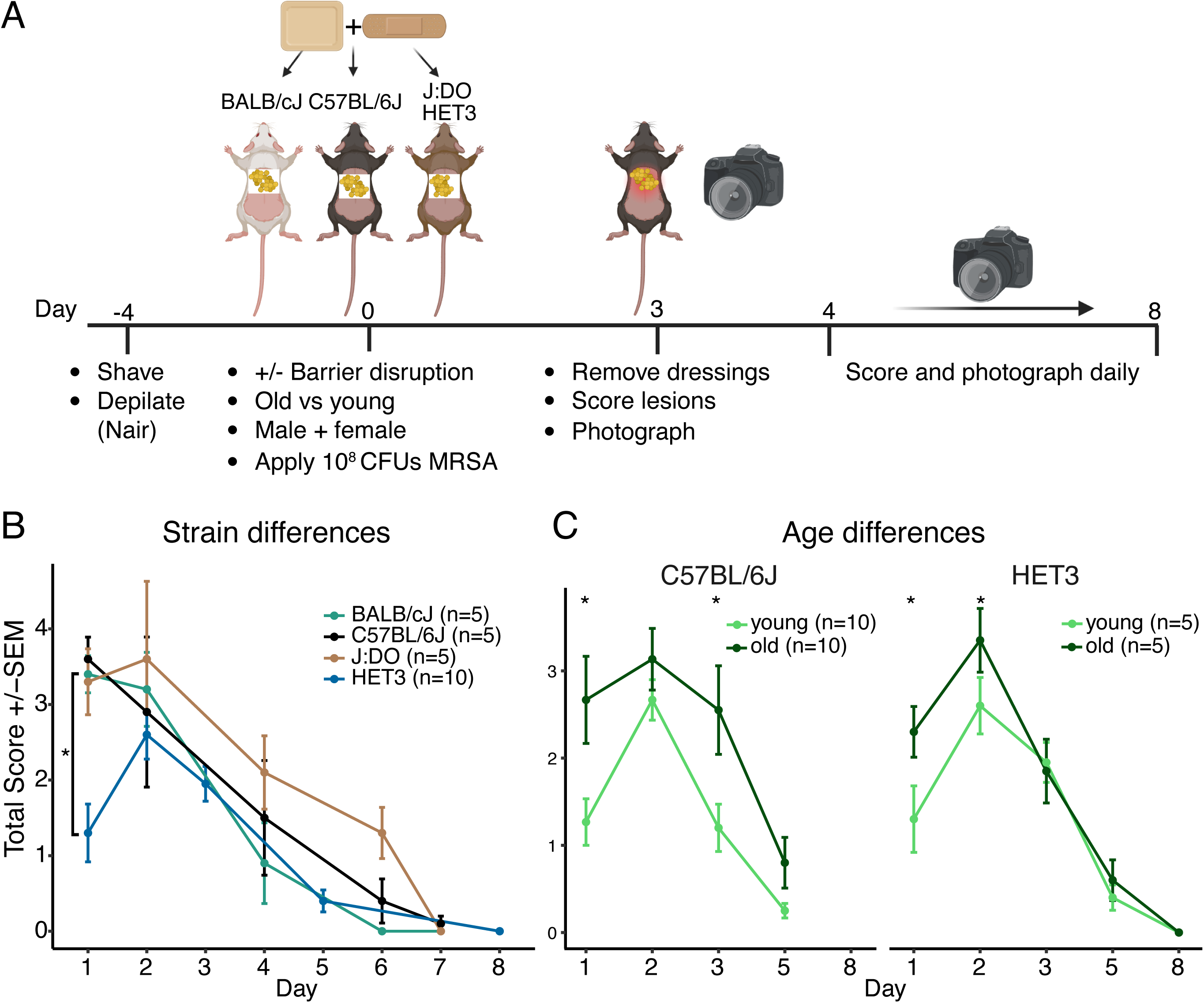
Study design, model establishment, and lesion dynamics in different mouse genetic backgrounds. (**A**) Experimental setup for *S. aureus* infection model. (**B**) Four mouse strains (C57BL/6J, BALB/cJ, J:DO (all n=5, females), HET3 (n=10, females) were tested for strain- specific differences in lesion development and resolution. The sum of all scores (scaliness, redness, skin breakdown) was used as a proxy for the overall phenotype development and plotted with the mean standard error (SEM), showed similar infection resolution kinetics between mouse strains. (**C**) Lesion resolution of young and old mice of an outbred versus inbred strain (HET3, n(young) = 5, n(old) = 5, C57BL/6J n(young) = 10, n(old) = 10) showed similar lesion resolution trajectories across strains and a. A linear mixed model was used to determine significance between groups to account for missing data points. *Statistically significant with p<0.05. ns: not significant, SEM: Standard Error of the Mean.

Although lesion scores varied across strains, the lesion resolution trajectory was largely comparable. HET3 mice exhibited delayed resolution relative to BALB/cJ (p=0.0194), but this difference was driven by day 1 post-exposure data and did not persist at later timepoints (**Figure 1B**). DO and HET3 mice showed a transient increase in lesion severity on day 2 following removal of the bacterial patch, before entering a recovery phase with convergent healing trajectories (**Figure 1B**). Similarly, healing trajectories were comparable across age groups (young: 8 weeks, or old: 80 weeks) in both C57BL/6J and HET3 mice, though aged mice tended to develop more severe lesions at early timepoints (**Figure 1C**). Middle-aged mice had variable phenotypes that did not suggest a linear relationship between age and infection outcome and were therefore excluded from subsequent experiments (**data not shown**).

We next examined whether barrier disruption or sex modified infection outcome. Because moderate barrier defects influence the outcome of *S. aureus* topical infection^16,17^, we tape-stripped the skin 10 times to partially remove the stratum corneum^18^ prior to infection in young and old C57BL/6J and HET3 mice (**Figure S1A)**. Tape-stripping modestly increased lesion severity on days 1 or 1 and 2 in most groups, but did not substantially alter overall kinetics. Histopathological analysis of infected skin showed no consistent differences between age or tape-stripping (**data not shown**). Sex had limited influence in this model, as healing kinetics were similar between males and females for the two parent inbred strains of the HET3 (BALB/cJ, C57BL/6J) (**Figure S1B)**.

To assess the global cutaneous response to infection, we performed bulk RNA-seq on infected and adjacent unaffected skin from the young and old C57BL/6J mice (**Figure S2A**). Principal component analysis (PCA) delineated age and infection groups (53% (PC1) and 29% (PC2) of the variance, respectively) (**Figure S2B**). Higher within-group variance was observed in old, infected animals (**Figure S2C**), consistent with increased heterogeneity in infection response in aging^19^. Both old and young animals had similar numbers of differentially expressed genes (**Figure S2D-F**), and shared upregulation of infection-related genes including *S100a8, S100a9*, *Klk6*, *Lcn2*, and *Mmp10*. Gene set enrichment analyses (GSEA) of hallmark genes revealed infection-associated differences, where old and young skin was enriched in terms related to immunity (i.e., complement, inflammatory response, and Jak-Stat signaling). (**Figure S2G**). CFU quantification also revealed no differences in *S. aureus* colonization levels between old and young animals (**Figure S6A**), supporting the similar responses observed at the transcriptional level.

Taken together, phenotypic and transcriptomic results suggest that while age and host variables may influence early lesion severity and inflammatory transcriptional state, ultimate infection resolution are similar in this model. Based on these findings, subsequent mechanistic experiments focused on young versus aged C57BL/6J mice for consistency and practicality.

### Microneedle profiling reveals age-dependent cutaneous immune dynamics after *S. aureus* infection

Our RNA-seq analysis showed a broadly inducible inflammatory response to *S. aureus* infection in both age groups (**Figure S2**). We next sought to capture the dynamic tissue resident cytokine and cellular response before and after infection with our microneedle patch (MNP) technology (**Figure 2A-B, Figure S3**). Alginate-coated MNPs enable minimally invasive, repeated sampling of skin interstitial fluid, allowing longitudinal assessment of cytokine secretion and immune cell infiltration without disrupting tissue integrity^20^.

**Figure 2:**
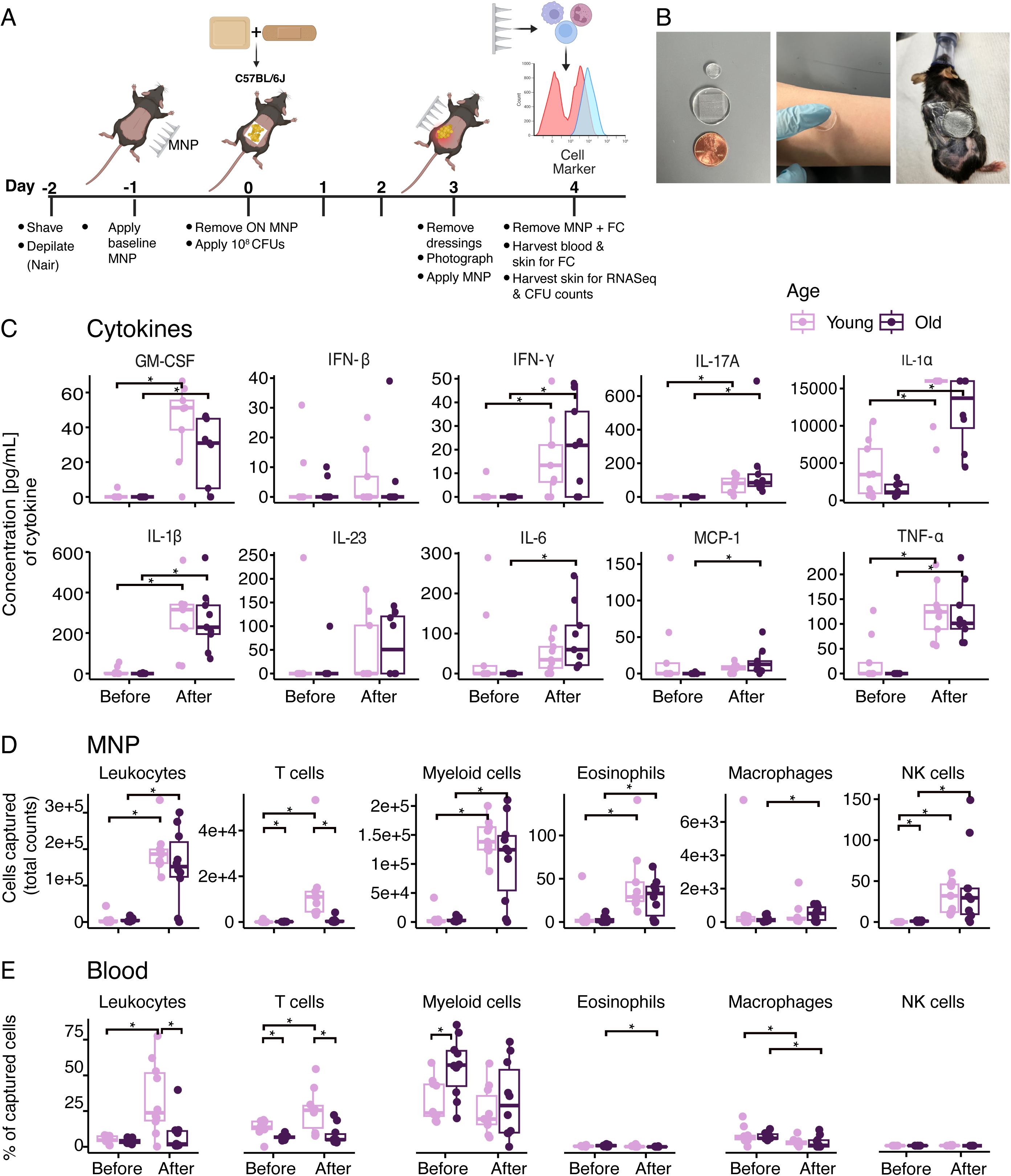
Local immune response captured by microneedle patches identifies age-associated differences in *S. aureus* skin infection response. **(A)** Experimental setup for *S. aureus* infection model and immune cell collection with young (8 weeks) and old (80 weeks) female (n=5) and male (n=4) C57BL/6J mice. For one young male, only data from blood were recovered due to spontaneous mortality. (**B**) Photograph of MNP relative to a penny, a human arm and a mouse. (**C**) Cytokine concentrations detected in interstitial fluid before and after infection, as measured by bead-based cytokine capture flow cytometry array showed both infection- and age-specific responses. (**D**) Immune cell counts detected in interstitial fluid before and after infection as measured by flow cytometry showed age-associated differences in T cell capture post-infection. d Paired Wilcoxon test was used to determine statistical significance between groups; *adjusted p- value (padj.) <0.05. MNP: microneedle patch, ON: overnight, FC: flow cytometry

We first used MNPs to assess the inflammatory cytokine milieu in skin pre- and post-infection. As anticipated for uninfected, intact skin, baseline cytokine concentrations and immune cell recovery were low across animals (**Figure 2C**). After infection, proinflammatory cytokines were significantly elevated in both age groups (**Figure 2C, Supplemental Table 1**), consistent with RNA-seq observations. Notably, IL-6 and MCP-1 levels were significantly increased only in old mice (padj. < 0.05), suggesting age-dependent enhancement of select inflammatory pathways after infection (**Figure 2C**).

We then compared MNP cell capture dynamics with conventional tissue dissociation methods on infected postmortem skin processed via enzymatic digestion and flow cytometry. Myeloid cells (∼60%) and macrophages (∼30%) were the predominant cell populations recovered in conventionally harvested skin in young and aged animals (**Figure S4A**). Conventional tissue harvests consistently yielded higher total cell numbers and greater animal-to-animal variance in aged animals compared to young, regardless of cell type (**Figure S4B**). By contrast, variance within groups was markedly lower in MNP-collected cells (**Figure 2D**), supporting the utility of MNPs for resolving tissue-level immune dynamics.

Leukocytes, myeloid cells, NK cells, and eosinophils increased significantly post infection regardless of age (**Figure 2D**, padj < 0.05). Notably, young mice showed a significant post-infection increase in T cells (p = 0.00052) not observed in aged mice, suggesting age-dependent differences in cutaneous T cell dynamics (**Figure 2D, Supplemental Table 2**), though these data do not differentiate altered T cell recruitment, retention, activation, or survival within infected skin.

Because tissue-resident versus circulating immune responses may be differentially affected by age, we also analyzed peripheral blood collected pre- and post-infection (**Figure 2E**). T cells, eosinophils, macrophages, NK cells, and leukocytes represented a small (<10%) fraction of total cells captured pre-infection (**Figure 2E)**. T cells and leukocytes accounted for a greater fraction of captured cells pre- and post-infection in young compared to old mice, trending towards significance. (**Figure S4CD, Supplemental Table 3**). These results suggest that the local immune cell dynamics captured by MNPs have dynamics distinct from those in circulation. Finally, sex-specific differences in cytokine and immune cell levels from skin tissue, blood and MNPs were negligible (**Supporting Data in Supplemental Material and Methods**), so for practicality we focused subsequent experiments on female mice for consistency.

We next used longitudinal MNP sampling to define how the tissue resident immune response evolved during the resolution of *S. aureus* infection over 7 days (**Figure 3A**). Despite substantial within-group heterogeneity in cytokine levels, major conclusions from the previous cross-sectional analyses were largely recapitulated. GM-CSF and IL-1β peaked one day after baseline measurement and declined rapidly in both young and aged animals (**Figure 3B**). In contrast, IL-17*a* peaked earlier in old animals (day 1) but was delayed until day 3 in young animals. Similarly, IFN-*γ* induction was observed only in young animals and also peaked at day 3, whereas levels remained low in aged mice. Other cytokines, including IL-1*a*, IL-6, MCP-1, and TNF-*a* peaked at day 3 in both age groups, with TNF-*a* trending higher in young mice. IL-23 and IFN-β levels remained low throughout the timecourse.

**Figure 3:**
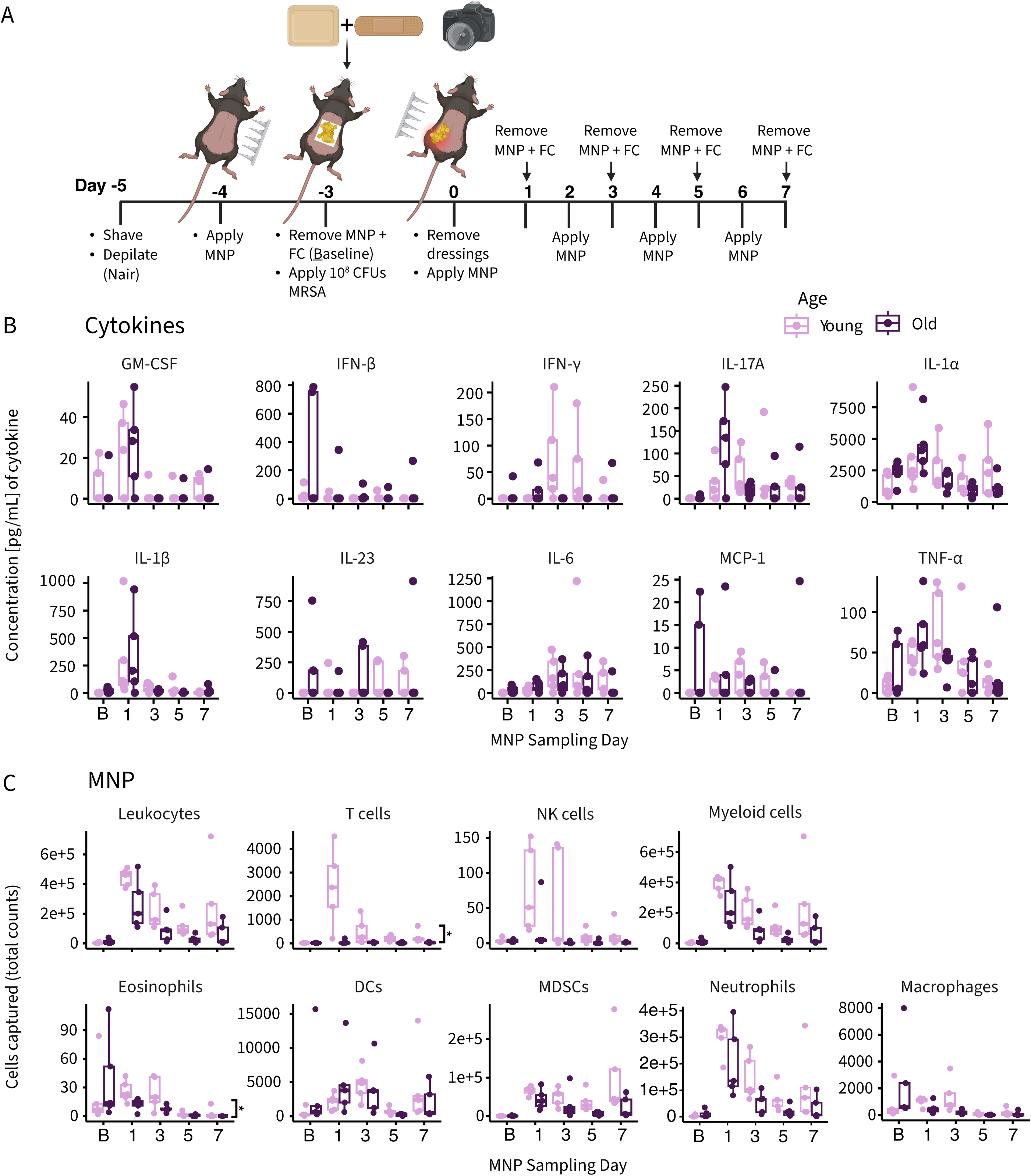
Longitudinal immune response captured by microneedle patches identifies altered immune dynamics in aged mice after *S. aureus* infection. **(A)** Experimental setup for *S. aureus* infection model including timepoints for cytokine and immune cell collection from young (8 weeks, n=5) and old (80 weeks, n=5) female C57BL/6J mice. (**B**) Cytokine concentrations detected in interstitial fluid before infection and during resolution as measured by bead-based cytokine capture flow cytometry array showed age- and time-associated trends in cytokine capture. (**C**) Immune cell counts detected in interstitial fluid sampled before infection and throughout infection resolution as measured by flow cytometry showed significant age-associated differences in T cell and eosinophil capture over time, with age-associated but non-significant trends in NK cells and neutrophils. Significance between groups was determined with a linear mixed model; * adjusted p-value (padj.) <0.05. MNP: microneedle patch, ON: overnight, B: baseline, FC: flow cytometry.

Immune cell recovery by MNP sampling was also time dependent. Recruitment of leukocytes, macrophages, and dendritic cells was highest on days 1 and 3 post-infection in both young and old mice (**Figure 3C**). Consistent with the single time point experiment, T cell numbers peaked in young mice at day 1 post infection, while numbers for aged mice remained low throughout the time course (**Figure 3C**). T cells also persisted longer in young vs. aged tissue, indicating differences in the rate of clearance over time. NK cells, neutrophils and eosinophils were also elevated in young mice on days 1 and 3, then declined during resolution. Notably, the rate of eosinophil clearance was significantly higher in aged animals compared to young, indicating age-associated divergence in both activation and resolution phases of the tissue immune response (**Figure 3C).**

Finally, analysis of circulating immune cells from peripheral blood (**Figure S5A**) indicated that T cell percentages rose gradually and peaked at day 5 in young mice, lagging behind their day 1 peak in the skin (**Figure 3C**). This temporal difference suggests that the early T cell response captured in skin may reflect activation or retention of tissue resident T cell populations, rather recruitment from circulation alone. As in our single timepoint experiment, circulating T cell percentages trended higher in young versus old mice, here regardless of timepoint. Myeloid cells were the predominant circulating cell population and declined over time as infection resolved irrespective of age. Few circulating macrophages, eosinophils, dendritic cells, or neutrophils were identified, consistent with the localized nature of the epicutaneous infection model.

Taken together, these data show that microneedle profiling captures age-associated immune dynamics at the site of infection that is not fully reflected in blood or bulk tissue measurements. Young mice mount an early cutaneous T cell response that precedes detectable systemic changes, whereas aged animals exhibit a blunted and temporally altered local response. More broadly, these findings indicate that aging reshapes the kinetics and magnitude of skin-localized immune activation during epicutaneous *S. aureus* infection.

### Bulk RNA-seq reveals largely conserved age-dependent skin responses to epicutaneous *S. aureus* infection

In parallel with single timepoint-based microneedle-based profiling (**Figure 2**), we characterized the paired cutaneous transcriptional responses to the epicutaneous infection to understand if and how MNP and transcriptional data correlate within the same animal. Infected and adjacent uninfected skin were collected at the peak of infection and processed for bulk RNA-seq (**Figure 4A**). Again, bacterial burden (CFUs) was similar between young and age tissue (**Figure S6A**), consistent with the early harvest time point prior to infection clearance and the ability of *S. aureus* to persist on occluded skin^12^, and was thus not analyzed as a covariate.

**Figure 4:**
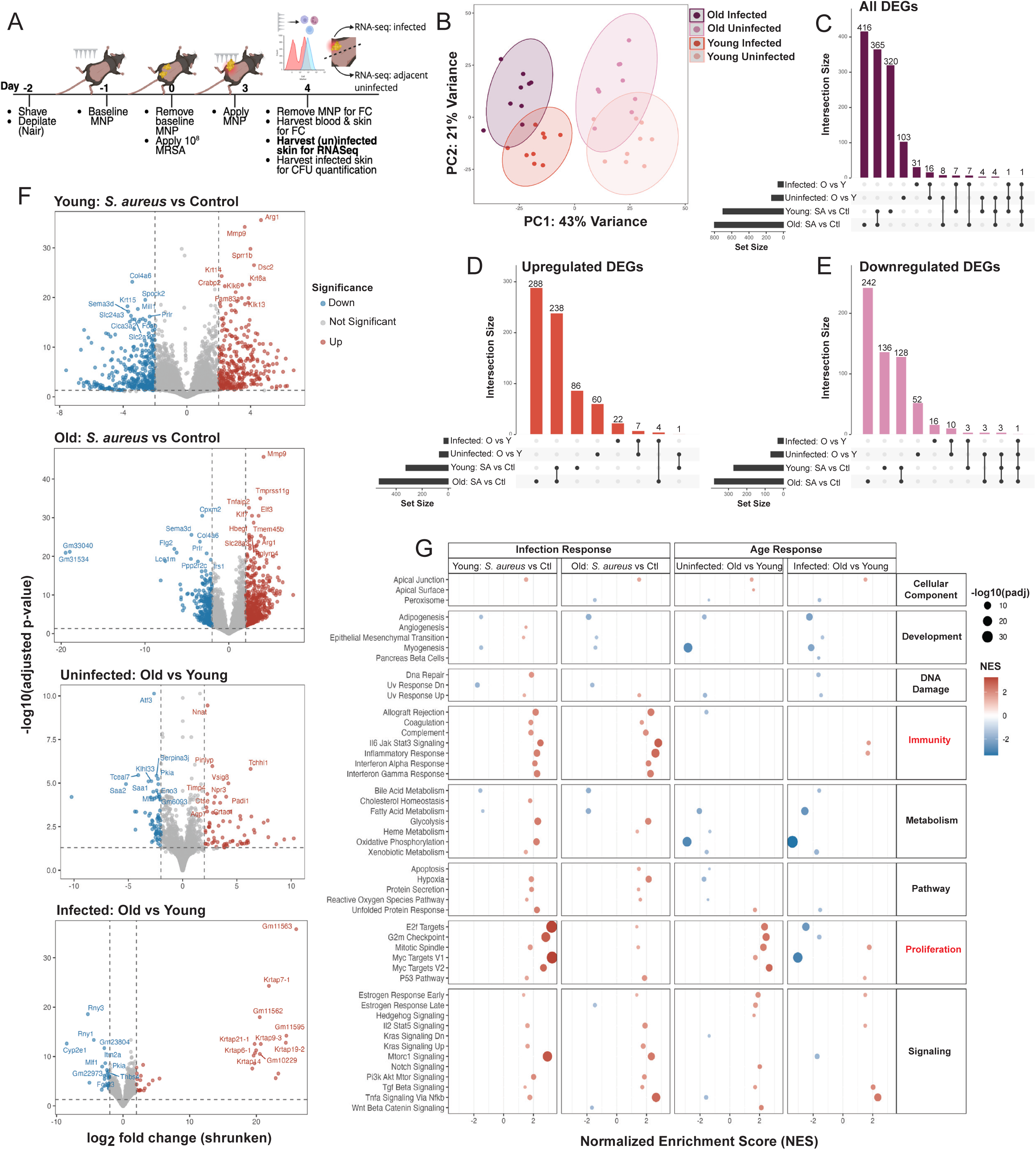
RNA-seq defines age-associated skin transcriptional programs in response to *S. aureus* infection. **(A)** Young (8 weeks) and old (80 weeks) male (n=9) and female (n=9) C57BL/6J mice were epicutaneously exposed to *S. aureus* (1×10^8^ CFU) for 3 days. MNPs were applied to sample ISF at the height of infection, then a 2cm^2^ full-thickness infected or adjacent uninfected skin biopsy was harvested. (**B**) Principal component analysis of variance normalized data for all comparison groups showed infection (PERMANOVA R^2^ = 0.20468, F = 10.45, p = 0.001) versus age (PERMANOVA R^2^ = 0.09939, F = 5.0738, p = 0.001) as a dominant driver of transcriptional response. (**C**) Upset plots of all differentially expressed genes (DEGs, |Log2FC ≥ 2| and padj. ≤ 0.05) (**D**) or subset by upregulated (**E**) or subset by downregulated gene sets showed largely shared gene sets between infection groups regardless of age or direction of gene expression. (**F**) Volcano plots of each comparison type with red points indicating significantly upregulated (Log2FC ≥ 2 and padj. ≤ 0.05) genes and blue points indicating significantly downregulated (Log2FC ≤ -2 and padj. ≤ 0.05) genes. Old and young infection groups shared many upregulated infection response genes (e.g., *Mmp9*, *Arg1*). In age-associated comparisons, old transcriptomes were dominated by upregulation of proliferation and keratin associated genes. (**G**) Gene Set Enrichment analysis using Hallmark gene sets (padj. < 0.05) for each comparison. The 5 most enriched or depleted processes (Normalized Enrichment Score, NES) showed shared and enriched infection-related processes between old and young infected tissue. Young tissues were more strongly enriched in proliferation and signaling associated terms. CFU: colony forming unit, MNP: microneedle patch, ISF: interstitial fluid, Log2FC: Log2 fold change.

Infection and age were the primary drivers of global transcriptional profiles with infection and age explaining 20.5% and ∼10% of variance, respectively (PERMANOVA R^2^ = 0.20468, F = 10.45, p = 0.001; R^2^ = 0.09939, F = 5.0738, p = 0.001). Differential expression analysis showed a largely conserved infection response between old and young infected tissues (**Figure 4C-F**). Both young and aged infected tissue showed robust induction of proinflammatory and antimicrobial genes, with some shared elements (e.g., *Nos2, Il1b, Cxcl2*, and *Defb3)*, but also age-specific responses. Genes uniquely upregulated in young infected tissue included *Spink12, Serpinb1b, Mmp1a, Il33, Il6, Cxcl1,* and *Klk8* (**Supplementary file 1**). Aged infected tissue had a unique, robust anti-infection induction of *IL-17f, Tnf, Lcn2, Mpo, Ly6g, Il-22,* and *Il-23α,* among others. No age-dependent transcriptional changes of cytokines differentially present in the MNP cytokine protein array were observed (**Figure S7**), suggesting that transcriptional changes at this timepoint may not capture the localized protein-level changes detected in the interstitial fluid. Aged and young skin shared infection responses in IL6 Jak Stat3 signaling, complement, and interferon response at the GSEA level (**Figure 4G**), including, surprisingly, proliferation-related terms including E2F targets, G2m checkpoint, and Myc targets. However, young infected skin proliferation gene sets were dominated by canonical inflammation-associated genes (e.g., *Il1a*, *Socs1*) and immunometabolic genes (e.g., *Pgk1*, *Ldha*), while old uninfected proliferation gene sets contained cytoskeletal genes (e.g., *Arhgef7*, *Trio*) and some senescence associated genes (e.g., *Pml*, *Cdkn2a*, *Nbn*), suggesting that older epithelial tissues may be less responsive to infection, while young tissues respond to infection through both immune and proliferative pathways.

Thus, while microneedle profiling revealed age-associated differences in tissue resident immune cell dynamics and cytokine abundances, bulk RNA-seq, which likely is predominated by the epithelial barrier response to infection, identified a broadly shared proinflammatory transcriptional response across age. These results suggest that generalized infection-responsive transcriptional programs in skin are largely intact in aging, which more strongly manifests at the level of local immune cell dynamics and interstitial cytokine responses.

### Ovalbumin sensitization reveals age- and microbiome-dependent skin immune dynamics distinct from *S. aureus* infection

Because *S. aureus* infection combines microbial, inflammatory, and barrier-associated cues, we next asked whether the age-dependent cutaneous immune dynamics observed after bacterial challenge reflected a generalized response to skin disruption or were shaped by the microbial context of the stimulus. To address this, we used an ovalbumin (OVA) sensitization model as a mechanistically distinct, noninfectious complementary comparator of localized allergic/barrier-driven inflammation^21,22^. Longitudinal MNP sampling revealed a time and location-dependent cellular response to OVA sensitization in which OVA-treated young mice tended to accumulate more leukocytes, myeloid cells, and macrophages in later stages of sensitization compared to aged mice (**Figure S8, S9**). Young mice also showed greater eosinophil recovery than aged animals, suggesting age-dependent differences in allergic inflammatory dynamics after OVA sensitization. Notably, MNPs were again essential to capturing these transient intermediate immune responses, as no differences between groups were observed at endpoint.

Because commensal microbes can also influence barrier immunity, we also incorporated antibiotic treatment into the OVA sensitization model to reduce background microbial influence and to facilitate interpretation of OVA-driven inflammation. This contrasts with the *S. aureus* infection model, where preserving the natural host–microbe interaction was central to the experimental question. Systemic and/or topical antibiotics (ABX) strikingly depleted skin and blood leukocytes and myeloid cells, and to a lesser extent, T cells (**Figure S10A-C**) irrespective of age (or dose). While this depletion was transient, ABX altered subsequent cellular immune dynamics, with ABX/OVA treated animals showing more similar response patterns across age groups for most cell types, a finding further supported by the ABX-only group. However, young ABX/OVA treated animals tended to mount a stronger macrophage, DC, and eosinophil response that was not matched in old animals, again suggesting differential sensitization dynamics. Finally, as with the *S. aureus* infection model, immune dynamics captured with microneedles were not fully mirrored systemically; peripheral blood profiling showed that most immune cell populations followed broadly similar longitudinal dynamics across groups and timepoints, albeit with high variance between groups (**Figure S9**).

Together, these results suggest that the OVA sensitization is distinct and localized, is not mirrored systemically, and is unique from the response associated with *S. aureus* infection. At the same time, systemic microbial depletion broadly reshaped the cutaneous immune environment and appeared to blunt several age-associated differences, indicating that microbiome depletion itself can modify local immune dynamics. Thus, these findings support a model in which aging shapes the skin immune responses in a stimulus-dependent manner, while microbial perturbation can further alter the tissue immune context versus systemic immune responses.

## DISCUSSION

Aging skin undergoes profound structural, epithelial, microbial, and immunologic remodeling, yet there are few mechanistic insights into how aging affects how the cutaneous immune system senses and responds to bacterial infections. Moreover, aging does not occur linearly, but in different trajectories that can be difficult to model with *in vivo* studies. Here, we characterized age-associated dynamics of immune and epithelial responses in a mouse skin infection model, including sex, and genetic background as variates. Through transcriptional profiling of skin together with longitudinal, microneedle patch sampling of immune cells and cytokines in the skin’s interstitial fluid, we defined age-associated signatures and dynamics in the cutaneous response to *S. aureus* infection.

Importantly, we found that although young and aged mice developed similar lesions and cleared infection with comparable kinetics, longitudinal immune sampling was key to revealing marked age-dependent differences in the timing and composition of the local response. These results suggest that similar infection outcomes may mask substantial age-dependent differences in the immune pathways used to achieve resolution. Here, young animals mounted an early and robust T cell response and stronger neutrophil and eosinophil recovery after infection, suggesting that aging alters both innate and adaptive immune dynamics even when infection outcomes are similar. While our study did not directly test reinfection or protective memory, this delayed and attenuated T cell response in aged animals raises the possibility that aging could impair the capacity of the skin to rapidly mobilize immune responses in subsequent microbial encounters.

Additional extrinsic or intrinsic factors not captured in our model may also be important in the aged skin response to infection. For example, we did not formally address the contributions of lifestyle or comorbidities, which may render aged skin more susceptible to a variety of insults or serious complications. Factors including diabetes, obesity, and/or frailty, as well as exposure to high risk environments like hospitals or long-term care facilities where *S. aureus* and other pathogens circulate^9^ are important future considerations. One metanalysis estimated that methicillin resistant *S. aureus* was present in at least 15% of residential care facilities globally^23^, suggesting broad baseline risk associated with aging care across the world. It is also of great interest to contrast infection outcomes in healthy versus unhealthily aged mice – the murine equivalent to human frail, or unhealthy aging, in which the skin microbiome shows the greatest dysbiosis^24^ and potentially cutaneous immune response. Future *in vivo*^25,26^ or *in vitro* models could incorporate comorbidities or other lifestyle risk factors to better parse their influence on infection risk and resolution in this population.

Notably, our MNP sampling allowed us to identify an early, strong T cell response to infection in young animals in this model that was largely absent in older animals. Although these data reveal clear age-associated differences in local immune dynamics, the underlying drivers remain to be defined. For example, our study does not distinguish whether these differences arise from altered tissue retention, recruitment from circulation, antigen presentation, epithelial signaling, or other age-associated changes in the local barrier microenvironment. Lagging changes in matched blood immune cell frequencies indicated that this was likely a tissue-specific response, rather than one mediated through circulating immune cells. Similarly, increases in neutrophil and eosinophil populations were identified only in young skin, suggesting an altered innate and adaptive cutaneous milieu in aged animals that may be decoupled from physical infection resolution. Furthermore, we identified the most age-dependent variability at the cellular, rather than the cytokine level, which were infection-driven but largely irrespective of age. These results suggest that local cutaneous immune cell populations may play dominant and age-variable roles in controlling epicutaneous S*. aureus* infection.

In addition to our phenotypic and cellular assessments, we also found age-associated transcriptional differences in the epithelial response to *S. aureus* infection. There are very few reports of age- and infection-associated transcriptional changes in mouse skin, with most aging studies focused on uninfected or mechanically damaged tissue^27–29^. In young mice, eosinophils contribute to skin inflammation in an IL-17A and IL-17F dependent manner post *S. aureus* infection^30^, and T cells^31^ and keratinocytes^32^ furthermore mediate the IL-17 response to epicutaneous *S. aureus* infection, demonstrating the complex interplay of innate and adaptive cutaneous immunology. Consistent with this, our transcriptomic and MNP analyses revealed that both young and old animals shared an IL-17α transcriptional and cytokine response to infection. We did not identify strong age-association in the expression of genes within the IL-17 pathway (i.e. *Mmp3*, *Tnf-α*, *Il-1β*, *Il-13*, *Il-6*, and *Ccl7*), although this could in part be due to the decoupling of protein from gene expression, or the inability of bulk RNA-seq to differentiate cell-type specific transcriptional patterning in low abundance cells like T-cells or eosinophils. Also consistent with previous reports, our MNP data revealed that young animals mounted strong T cell and eosinophilic^30^ responses to *S. aureus* infection. The blunted T cell and eosinophil response in aged animals may reflect impaired antigen presentation or altered cytokine milieu, which will be mechanistically dissected in future studies.

To determine whether age-associated immune dynamics were specific to bacterial infection or reflected a broader response to challenge, we examined a complementary OVA sensitization model. OVA sensitization also elicited a localized immune response that differed from the *S. aureus* response in both timing and cellular composition. This suggests that aging skin immune dynamics are not fixed, but rather dynamically shaped by the nature of the local stimulus and the surrounding microbial context. Indeed, antibiotic treatment itself broadly alters cutaneous immune cell recovery, and when taken together with OVA, the antibiotics blunted several age-associated differences. This suggests that commensal-dependent cues may contribute to how young and aged skin respond to barrier challenge. We note that these data should be interpreted primarily as evidence that barrier sensitization responses are shaped by both age and microbial context, rather than as a direct one-to-one comparison with the infection model, where we did not use antibiotics to preserve the native infection context and capture the full host response to bacterial exposure.

To our knowledge, this is the first report to use an epicutaneous infection model to understand host genetic factors that may influence infection outcome, and as a function of age. Previous studies have demonstrated that mouse genotype can contribute to infection outcome in more severe models of infection^33^. Outbred mice had significant genetic determinants of infection resistance, where certain chromosomal loci in Collaborative Cross mice were important for early versus late susceptibility to systemic *S. aureus* infection^34^. Although we did not identify phenotypic differences in the severity or resolution of *S. aureus* infection across the mouse strains we tested, this may be due to the nature of our model, in which we investigated a tissue resident response without dissemination of infection from the skin to other organs. Additionally, for practical constraints, we did not use every combination of strain (aged BALB/cJ or DO strains are not readily available), age, and sex in our experiments which could reveal infection response nuances not captured in this study.

While we did not detect strong sex-dependent differences in our phenotypic outcomes, there is evidence to suggest sexual dimorphism in the response to *S. aureus* infection. Both human and murine males are more susceptible to *S. aureus* infection, while female mice were shown to be protected from dermonecrotic damage in a model of SSTI^35,36^. We did not identify significantly different burdens of *S. aureus* between males and females, nor did we find significantly different immune cell profiles in males versus females at our initial harvest time point. However, both our infection model and CFU sampling technique differed from these previous reports of sex differences in the epicutaneous model^37^. Future studies could include both males and females and additional clinically relevant isolates of *S. aureus* and other skin pathogens such as *Streptococcus pyogenes*^38^ to further resolve sex- or genotype-dependent responses to infection.^33,39,40^

Importantly, this work established the feasibility and power of MNPs for repeated, non-destructive sampling of skin immune responses in live animals. Unlike traditional terminal approaches, MNPs preserve spatial and temporal context and can capture real-time immune changes across the course of infection, with an animal serving as its own baseline control. This platform represents a valuable tool for continued investigations into age- and immune-related vulnerabilities in skin infections and is well-validated in mouse and human models of vaccination^13^ and skin inflammation^20^. However, logistical and biological challenges remain for MNP sampling in animal models. For example, repeated anesthesia exposure is required to place MNPs, which can be stressful or cause spontaneous mortality particularly in aged animals. Repeated band aid application and removal required for longitudinal MNPs studies is also stressful to skin tissue and can damage adjacent skin areas surrounding the MNPs. In addition, we did not discern whether MNP application exposes the dermal layer to microbes, potentially altering infection response. Further, while we included baseline measurements for every mouse we did not include a group that received vehicle administration followed by longitudinal MNP sampling, which would be informative in understanding broader impacts of MNP application on skin biology and barrier integrity. Work is ongoing to understand how MNP application may influence resident skin microbiota and skin barrier function. Nonetheless, MNPs are a powerful tool for cutaneous cell sampling in animal models, with exciting future applications not only in functional characterization but also in aging biomarker discovery and drug delivery^41^.

In summary, our phenotypic, transcriptional, and longitudinal immunophenotyping analyses support a model in which aging may not merely weaken skin immunity, but reshapes the timing, cellular composition, and microbial dependence of local immune responses to challenge. Our study, which leverages a powerful tool for skin profiling, lays the groundwork for functional characterization of immune cell and epithelial changes in aging, and more broadly across the human lifespan.

## MATERIALS AND METHODS

### Animals

C57BL/6J (Strain code: 000664), BALB/cJ (Strain code: 000651), J:DO (Strain code: 009376) and (CByB6F1/J x C3D2F1/J)/J (from Strain code: 036603, here on abbreviated as HET3; also known as UM-HET3) mice were obtained from The Jackson Laboratory, Bar Harbor, ME, USA. Young mice were 6-9 weeks, middle-aged mice 33 weeks, and old mice 80-83 weeks old at experimental start.

All mice were housed in the animal facility at the University of Connecticut Health Center (UCHC) in Farmington, CT, USA. All experiments involving mice followed protocols that were approved by the UCHC Institutional Animal Care and Use Committees (IACUC) and were performed in accordance with National Institutes of Health regulations. Five mice were co-housed and maintained under standard day-night cycles. At the endpoint, mice were euthanized by CO2 asphyxiation.

### Bacteria

*Staphylococcus aureus* USA300 LAC (USA300-0114, NR-46070, BEI) was streaked on blood agar, and then a single colony was inoculated into tryptic soy broth (TSB, Fisher Scientific, DF0370-17-3) and grown overnight at 37°C with shaking at 200 rpm.

Colony forming units (CFUs) per 1 ml per 1 OD were assessed using hemocytometers (Fisher Scientific, 22-600-107) three times and then used for subsequent experiments (1OD equals ∼ 2×10^8^ CFU). For microbial application, the overnight culture was refreshed at 1:200 and grown to the exponential phase (∼OD = 1) in TSB at 37°C and shaking at 200 rpm. Cells were washed once by pelleting and resuspending in sterile PBS (phosphate-buffered saline). After a second centrifugation, the supernatant was decanted, and the pellet resuspended and diluted in PBS to obtain a CFU count of 10^8 CFUs per 150μl.

### Skin barrier disruption

3-5 days prior to the experiment, mice were anesthetized with 2-5% isoflurane in oxygen, placed on a heating pad (37°C), and eyes were lubricated with ointment (OptixCare). The mouse dorsum was shaved and depilated with Nair (Nair Hair Removal Cream). On experimental day 0, mice were anesthetized as described above. A piece of tape (Shipping Tape, 3M) of ca. 10 cm was loosely wrapped around 2 fingers and taped together to form a ring. The mouse’s dorsal skin was gently stretched and the tape placed (with the 2 fingers inside the ring) on the skin and removed in one fast motion. Repeating this procedure 10 times has been shown to weaken the skin barrier using transepidermal water loss (TEWL) measurements using a Tewameter (Courage + Khazaka electronic GmbH)^16,18^. Lower TEWL values are indictive for intact skin and higher values for decreased skin barrier function.

### *S. aureus* infection model

To model *S. aureus* epicutaneous infection, we applied *S. aureus* in high density onto dorsal mouse skin for 3 days as previously described^31,42^. The dorsal fur was shaved and depilated (Nair Hair Removal Cream) at least 2 days before further treatment. On experimental day 0, mice were anesthetized (as described above). A sterile gauze pad (1.5cm x 1.5cm, CVS Health Sterile Non-Stick Pads) was soaked with 150 -255 μl *S. aureus* solution containing 10^8^ CFUs in PBS and applied to the dorsal skin of the mouse. The pad was secured with breathable bioocclusive dressing (Tegaderm, 3M) and wrapped with 2 layers of adhesive band-aids to secure the Tegaderm. Mice were monitored daily and the outer band aid replaced if necessary. After 72 hours, mice were anesthetized and the bandaids and gauze pad were gently removed.

### Skin lesion assessment

Mice were briefly anesthetized for lesion assessment. We scored redness (none (0), slight (0.5- 1.5), moderate (2-2.5), severe (3)), scaliness (none (0), slight (0.5-1.5), moderate (2-2.5), severe (3)) and skin broken (no (0), yes (1)) and then compared the sum of all parameters (Total score) between mouse groups. Lesions were assessed daily or every other day up to 8 days after removal of the *S. aureus* pad. Pictures were taken to document the progress.

### Skin sample collection

Skin tissue – approximately 1 cm^2^ of affected (*S. aureus* induced lesions) and unaffected (macroscopically healthy adjacent tissue) - was collected postmortem for RNA extraction, CFU enumeration, and flow cytometry.

For RNA extraction, the tissue was placed in RNALater (AM7024, Invitrogen), kept at +4°C overnight and then frozen until RNA extraction. For RNA-seq, the epidermis was not separated from the dermis as extensive pilot experiments did not reproducibly result in high quality RNA for epidermis or dermis (data not shown). Varying length of digest time with Dispase II (D4693, Sigma-Aldrich) (15 min to 4h at 37°C, and overnight at 4°C, 2.5 U/mL in 1x PBS) and different skin pretreatment methods (RNALater, flash freezing, fresh) were evaluated.

For CFU enumeration, approximately 1 cm^2^ of infected tissue was excised, weighed, and homogenized in 500uL of PBS with 1mm silica/zirconia beads (Biospec cat# 11079110zx). Each sample was serially diluted, plated on blood agar plates, and incubated overnight at 37°C. For fluorescence-activated cell sorting (FACS), the skin was minced and processed as described below.

For histological evaluation, affected with adjacent normal-appearing skin of young and old mice (tape-stripped and not taped) was collected and fixed in 4% PFA (+4°C) overnight. Tissue was washed in 1x PBS, color coded and stored in 70% Ethanol as described below.

### Histology

Fixed tissues were gently washed twice in PBS and then color-coded using tissues dyes (Individual Davidson Marking System® dyes) to embed multiple tissues per FFPE block. Color-coded tissues were stored in histology cassettes in 70% ethanol until paraffin-embedding, sectioning (5 μm) and H&E staining using standard protocols at the Histopathology facility at The Jackson Laboratory, Bar Harbor, ME.

Lesions were scored blindly by a histopathologist. For the epidermal measurements, viable cells (basement membrane to top of granular layer) then total (first measurement plus attached stratum corneum) were measured in five different fields of view (excluding lesions such as ulcers or crusts). Ulcers are lesions that extend through the basement membrane. Crusts are desiccated serum and dead inflammatory cells from the ulcers or ulcers lateral to the section. Hypodermal fibrosis is an indication of severe ulcers that have started to or have healed.

In more detail, epidermal thickness for the Malpighian layer (live cells) and full thickness (subtracting Malpighian layer from full thickness results in the stratum corneum thickness) were measured for areas without crusts or ulcers at 40x magnification. Percentage of scabs and ulcers, as well as counts of ulcers and hypodermal fibrosis were recorded in each sample. To assess group differences, the non-parametric (unpaired) Wilcoxon signed-rank test was employed for matched (but independent) comparisons. P-values were post-hoc multiple-test corrected using the false discovery rate (FDR) method. Corrected p-values (padj.) <0.05 were deemed significant.

### Blood collection and isolation of PBMCs

Blood was collected for flow cytometry as baseline at the beginning of the experiment, after *S. aureus* infection, and at the endpoint. Blood was collected retro-orbitally with a hemato-clad capillary tube (21-176-6, Drummond) while the mouse was anesthetized. The blood was transferred to a vacutainer tube (K2EDTA, BD367856, BD Biosciences), rotated to mix the blood with the K2EDTA and kept at RT until the blood was mixed with 10x the volume of 1X RBC lysis buffer. After incubation at RT for 4-5 minutes with occasional shaking, the reaction was stopped by adding 3x the volume of 1X PBS. The cells were washed by spinning at 300-400 x g for 5 minutes at 2-8°C. The RBC lysis was performed 2 times, and the cells were resuspended in FACS buffer and transferred to a U bottomed 96 well plate for flow cytometry analysis.

### Dissociation of skin tissue to isolate immune cells

The weight of the skin tissue was measured before processing. Subcutaneous fat was removed from the dissected skin (∼ 0.5×0.5cm) and minced. The skin fragments were transferred to a 5 ml centrifuge tube containing 1 ml of digestion buffer, composed of 2mg/ml collagenase XI (Sigma, # C9407-1G), 0.5 mg/ml hyaluronidase (Sigma, # H3506-1G) and 0.1 mg/ml DNase (# DN25- 10MG) in RPMI supplemented with HEPES and Penicillin/Streptomycin. The tissue was further minced in the digestion cocktail using sterile scissors and 2ml of digestion cocktail was added. The samples were incubated in a shaker at 37°C for 45 minutes at 255 RPM. Following digestion, the homogenate was passed through a 100μm cell strainer into a 50 ml conical tube containing RPMI supplemented with 10% FBS and Penicillin/Streptomycin. The cell suspension was then centrifuged at 1500 rpm for 5 min at 37°C and the resulting cell pellet was resuspended and transferred to a U bottomed 96 well plate for flowcytometry analysis. The cell numbers were calculated per 100 mg of skin tissue collected.

### Fabrication of the Microneedle (MN) patches

Each MNP application yields approximately 30 μl of ISF and about 100,000 live cells demonstrating robust dual-phase temporal resolution: acute collection of cytokine fluxes within 20 minutes and immune cell profiling after a 6-18-hour application period^20^.

Microneedle (MN) patches were used to extract cytokines and immune cells from the skin of healthy and affected mice. MN patches are circular polymer patches (18 mm diameter) composed of an array of 400 tiny pyramidal needles that are 600 microns in height. The MN patches were fabricated in house by vacuum casting PLLA polymer at 200°C for 2 hours. The cooled MN patches were removed from the molds and sterilized by soaking in 70% ethanol overnight. The sterile, dried patches were oxygen plasma treated for 30 seconds to enhance the hydrophilicity of the patches and coated with alginate hydrogel Pronova SLG20 (Novamatrix #4202001) solution containing sucrose and dried overnight. The alginate hydrogel is crosslinked by using 20 mM calcium chloride solution and dried overnight. The prepared MN patches were stored in a desiccator until use. The materials used in the fabrication of the MN patches are FDA-approved and classified as Generally Recognized As Safe (GRAS) materials to be used on human and animal skin.

### Cytokine and immune cell sampling using microneedle (MN) patches

The MN patches were applied on the shaved skin of mice after anesthetizing with isoflurane. The patches were pressed gently on the skin and held for 10 seconds and secured using Tegaderm and band aids. To sample cytokines, the patches were left on the skin for 20 min, for immune cells, the patches were left on overnight and additionally secured using electrical tape. Electrical tape has enough strength to not be manipulated by other co-housed mice. The animals are closely monitored after application for signs of suffocation and once daily. For removal, the mice were briefly anesthetized, and the tape was cut off and removed. The sampled MN patches were placed on a 12-well plate containing the elution buffer (100 mM EDTA for cytokines and 100 mM EDTA supplemented with 10% FBS for immune cells).

The plates were placed on a shaker for 20 minutes to de-crosslink the alginate. After shaking the MN patches were held against the walls of the well plate using sterile forceps and washed thoroughly using the elution buffer in the well. For cytokine sampling, the eluent was collected in microfuge tubes and snap frozen by placing on dry ice containing ethanol and stored at -80°C for further cytokine analysis. The eluent containing immune cells were transferred to a U bottomed 96 well plate for flow cytometry analysis.

### Flow cytometry analysis of immune cells from the MN patches, blood and skin

The 96 well plate containing cells from blood, MN patches and skin tissue was centrifuged at 1200 rpm for 5 mins to pellet the cells. The supernatant was removed by aspiration and the pellets were resuspended in 100 μl of fixable Live/Dead Zombie NIR (Biolegend Cat #423106) (1:1000 in PBS) and incubated at room temperature for 15 minutes in dark. After incubation the cells were washed with FACS buffer by centrifugation at 1500 rpm for 5 minutes. The pellets were resuspended in 100 μl of anti-Fc receptor antibody (clone 2.4G2) and stained with surface antibodies in PBS and 2% fetal calf serum for 30 minutes on ice in dark. The cells were washed with FACS buffer after incubation and resuspended in 200 μl FACS buffer for flow cytometry analysis. All flow cytometry analysis was conducted on Symphony A5 (BD Biosciences), data was collected in Diva (BD Biosciences) and analyzed on FlowJo (BD Biosciences). The Antibodies used in these experiments are: CD45 (BUV395, Cat#564279, BD Biosciences), CD3ɛ (AF488, Cat# 100321, BioLegend), CD11b (BV785, Cat# 101243, Biolegend), NK1.1 (BV421, Cat# 108732, Biolegend), F4/80 (PE, Cat# 123110, Biolegend), CD193 (APC, Cat# 144512, Biolegend), CD11c (BUV661, Cat#750482, BD Biosciences), Ly6C (RB705, Cat#570264, BD Biosciences), MHC-II (BUV496, Cat#750281, BD Biosciences), and Ly6G (BUV805, Cat#741994, BD Biosciences).

### Cytokine analysis

The levels of inflammatory cytokines and chemokines were evaluated by a bead-based multiplex assay using the LEGENDPlex mouse inflammation panel kit (Cat#740150,740446 Lot#B445258 & B451622, Biolegend). The previously frozen eluents from MN patches containing the cytokines and chemokines were thawed on ice. After thawing the samples were mixed well by pipetting 2-3 times without formation of air bubbles. Undiluted samples were transferred to V-bottom plates in duplicate, and the assay was performed as per manufacturer’s instructions. The cytokine and chemokine concentrations were calculated using Biolegend’s LEGENDPlex data analysis software. The mean value of the duplicates was taken for analysis. In cases where one value was in range and the other above the detection threshold, we omitted the out-of-range value and duplicated the in-range-value. If both values were out of range, we kept both. Values that were below the detection limit were set to 0. For log2 fold change calculations of medians between groups, a pseudo-count of 1 was added to enable comparison of the groups. IL-10, IL-27, and IL12-p70 were include in the kit but excluded from further analysis as no induction was measured in any condition.

### Analysis and Statistics for skin lesions and FACS data

R (version 4.4.2) and RStudio (version 2024.12.1) were used for statistical analysis and data visualization. Plots were prepared with Rstudio packages tidyverse^43^, ggplot2^44^, ggpubr^45^, and reshape^46^.

The total scores of the lesions were compared between groups of interest over time using a linear mixed model (lmer, lme4 R package)^47^. Total score was used as the dependent variable, the interaction of the fixed effects (sex, age or tape stripping treatment) and day and the random effect accounting for the different mice (1|MouseID). Results were confirmed with the more robust and flexible Generalized Linear Mixed Models using Template Model Builder (glmmTMB, glmmTMB R package)^48^. emmeans function (emmeans R package) was used to perform post-hoc analysis on the fitted models^49^. p-values were post-hoc multiple-test corrected using the default Tukey’s method. P-values <0.05 were deemed significant.

Regarding the experiment assessing strain differences, original data was visualized (Figure 1B) but due to small differences in the timing of the HET3 observations, its data was shifted to allow for statistical strain to strain comparison as described above, i.e. Day8 was redefined as Day7, Day5 as Day6, and Day3 as Day4.

To calculate the rate of lesion healing, we performed a linear mixed-effects model analysis (lme4 package)^47^. The model included fixed effects for day, strain, and their interaction, with a random intercept for the individual mice to account for repeated measurements. The model estimates trajectories by fitting a linear relationship between lesion scores and time for each strain, while accounting for the correlation structure of repeated measurements from the same mouse. The lesion healing rates (slopes) were calculated as the estimated change in lesion score per day for each mouse group of interest using estimated marginal trends (emtrends, emmeans R package). These trends are computed by taking the first derivative of the model’s predicted values with respect to time, providing the rate of change in lesion scores. Pairwise comparisons of healing rates (slopes) between groups were conducted using the pairs function from emtrends, estimated marginal trends with the default Tukey’s adjustment.

To assess differences in immune cell and cytokine levels, we employed linear models accounting for the experimental design. For baseline and post-treatment comparisons, where each mouse contributed a single measurement per cell type or cytokine, we used linear models (lm()) with sex and age as fixed effects. To test whether sex differences varied between young and old mice, or whether treatment responses differed by sex, we included interaction terms in the models. For before-versus-after comparisons within age groups, where each mouse was measured at both timepoints, we used linear mixed-effects models (lmer(), lmerTest R package) with mouse as a random effect to account for repeated measures. For the longitudinal experiment, we excluded baseline cell counts from significance calculations as we were only interested in cell count trajectories in the context of lesion resolution. All analyses were stratified by cell type or cytokine and, where appropriate, by age group to examine sex differences separately in young and old mice. When deemed insignificant, sex was excluded from the model and data from male and female mice were pooled (Figure 2CDE, Suppl.Figure 3).

### Skin Sensitization and Microbiome Depletion Treatments

36 C57BL/6J mice were obtained from the Jackson Laboratory and houses under standard conditions with 12h light cycles. 18 mice were young (8 weeks) and 18 were aged (73 weeks). Mice were split in 4 groups. For 3 groups, 5 old and 5 young mice were used each. For Group 4, 3 young and 3 old mice were used as untreated controls to account for natural variability. Group 1 received oral antibiotics and the vehicle control topically (no ovalbumin). Group 2 received ovalbumin and oral antibiotics, while Group 3 received ovalbumin but was gavaged the vehicle control (no antibiotics)

### Antibiotic treatment

Mice were weighed before every round of antibiotics. Antibiotics were gavaged with 10ul/1g body weight at a final concentration of 10 mg/mL Ampicillin Sodium Salt (A0166-5G, Sigma-Aldrich), 10 mg/mL Neomycin trisulfate salt hydrate (N-620-5, Goldbio), 10 mg/mL Metronidazole (M1547-5G, Sigma-Aldrich) and 5 mg/mL Vancomycin hydrochloride (V2002-1G, Sigma-Aldrich). Stock solutions were prepared for all but Ampicillin. Ampicillin was dissolved freshly on the day of experiment and mixed with the other antibiotics. Metrodinazole was dissolved in DMSO, while the remaining antibiotics were prepared in sterile distilled water. The metrodinazole stock solution was prepared at 10x in order to obtain a final 10% DMSO concentration. The cocktail was sterile filtered with 0.22um filters. 10% DMSO were used as vehicle control. Antibiotics/Vehicle administration started 2 days before ovalbumin exposure and continued every day for a total of five times.

### Skin sensitization

To disrupt the skin barrier, we tape stripped each mouse 6x 24h before the administration of protein allergen (ovalbumin) to induce Atopic Dermatitis (AD) in young and aged mice^50^. Gauze (Sterile Gauze Pads, CVS Health or 2 layers of non-woven dental gauze (BRITEDENT)) of 1.6×1.6cm was moistened with ∼200ul of 1 ug/ul. The gauze was placed on the lower flank, covered with Tegaderm (3M) and then with a band aid followed by a layer of electrical tape or 3M Micropore Surgical Tape and left for 7 days. After 2 weeks break, the 7-day ovalbumin exposure was repeated. Mouse skin was exposed to ovalbumin for a total of 3 times.

### Microneedle application

Microneedles patches were applied overnight to collect immune cells before and after every ovalbumin application. For cytokine collection, microneedle patches were applied for 20 min after each Ovalbumin exposure. At baseline and endpoint, immune cells were sampled, processed for FACS immediately, and analyzed as described.

### RNA extraction and RNA-Seq of skin tissues

All RNA extraction and sequencing library preparation steps were performed in a sterile tissue culture hood. The skin was removed from RNALater, gently washed in PBS to remove excess salts and the adipose tissue was removed using a sterile scalpel. To increase RNA yield, skin was minced and placed into RLT-BME lysis buffer (1:100 BME (Beta-mercaptoethanol) (RNeasy 96 QIAcube HT kit) with two sizes of silica beads (50μl glass beads (0.1 diameter (BioSpec Products, 11079101) and 1 mm zirconia/silica beads (Biospec, Cat#11079110ZX) and physically disrupted for 3 minutes (30 Hz) using the TissueLyzer II (Qiagen).

RNA was extracted using RNeasy 96 QIAcube HT kit (Qiagen, Hilden, Germany) according to the manufacturer’s directions. Samples were eluted in nuclease-free water (Qiagen, Hilden, Germany) and frozen at -80°C until sequencing preparation. The RNA quality was evaluated using the 4200 TapeStation System (Agilent Technologies, Santa Clara, CA) with the High Sensitivity RNA ScreenTape Analysis (Agilent Technologies, Santa Clara, CA) or RNA ScreenTape Analysis (Agilent Technologies, Santa Clara, CA). RNA quantity was measured on the Qubit 2.0 Fluorometer (Thermo Fisher Scientific, Waltham, Massachusetts, USA). Sequencing libraries were prepared by combining the NEBNext rRNA Depletion Kit v2 (Human/Mouse/Rat) with NEBNext rRNA Depletion Kit v2 (Bacteria) (New England Biolabs, Ipswich, MA) and NEBNext Ultra II Directional RNA Library Prep Kit for Illumina (New England Biolabs, Ipswich, MA) following the manufacturer’s directions. Library quality was evaluated using the 4200 TapeStation System (Agilent Technologies, Santa Clara, CA) with the High Sensitivity D1000 ScreenTape Assay (Agilent Technologies, Santa Clara, CA). Library quantity was measured on the Qubit 2.0 Fluorometer (Thermo Fisher Scientific, Waltham, Massachusetts, USA). Samples were sequenced using Illumina NovaSeq. Reads per sample ranged from 16 million to 47 million with a mean of 27 million reads.

### Transcriptional profiling, gene ontology analysis

Raw RNA-seq data was processed with trimmomatic 0.39 to remove low quality reads^51^. Quality controlled reads were mapped to GRCm39 referece genome with STAR 2.7.1a^52^. Raw read counts aligned to each gene were computed with featureCounts v1.6.4 (from Subread 1.6.4)^53^. Raw read counts were normalized with RUVg^54^ using the 5000 most stably expressed genes across all samples. From the normalized counts, differential gene expression and principal component analysis was performed with DESeq2 (package version 1.44.0) using standard parameters (design ∼ age * infection)^55^. PERMANOVA was performed with the vegan R package version 2.7.3^56^. For volcano plot visualization, DEGs were normalized with the DESeq2 fold change shrink function and plotted with R package pheatmap version 1.0.12^57^. Shared and unique DEGs were visualized with R package UpSetR version 1.4.0^58^. Gene set enrichment analysis with Hallmark gene sets was performed using clusterProfiler^59,60^ (package version 4.12.6) on DEGs with a padj. ≤ 0.05.

## SUPPORTING DATA

### Histological observation

Histologic observation revealed high variability and a tendency towards a more severe phenotype in taped mice (as expected) but no significant difference between all combinations of old and young, tape-stripped and untaped mice (data not shown). Comparing all taped/untaped mice irrespective of age was equally not significant. Parameters assessed were full thickness, thickness of the Malpighian layer and stratum corneum, numbers of scabs, ulcers and hypodermal fibrosis and percentage of ulcers/scabs covering the tissue (Material and Methods). We refrained from further histological analyses.

### TEWL measurements and tape-stripping

We attempted to standardize the skin barrier effect caused by tape-stripping by repeatedly measuring the transepidermal water loss (TEWL) measurements as we tape-stripped the animals. However, measurements were variable and not linear with more tape stripping. We elected to tape-strip each mouse 10x (Material and Methods) which did result in increased TEWL value although not to a standardized value across the cohort.

### Assessment of sex-differences in cytokine and immune cell levels

The first experiment (Figure 2) included males and females in order to assess sex differences in immune and cytokine levels captured with MNPs, in blood or skin tissue. Statistical analysis (Material and Methods) showed no or minor significant differences for blood and MNPs, while the skin revealed statistical significances between female and male mice. Sex was included as a factor when assessing skin tissue (Figure S4CD).

However, skin was not collected in subsequent experiments due to the longitudinal nature (healing of the skin) of the experiments.

## Abbreviations

SSTI: skin and soft tissue infection;
DO: diversity outbred,
MRSA: Methicillin-resistant *S. aureus*,
MNP: Microneedle Patch,
CFU: Colony-forming Unit,
FC: Flow cytometry,
RNA-Seq: RNA sequencing

## DATA AVAILABILITY

The dataset related to this article can be found at the SRA under the BioProject accession number PRJNA1515871.

## ACKNOWLEDGEMENTS AND FUNDING

We are thankful to the Oh group for inspiring discussions and acknowledge the contribution of the Animal Care Team, Genome Technologies Service, Flow Cytometry Service and Histopathology facility at The Jackson Laboratory for expert assistance for the work described in this publication. We are grateful to Leonard M. Milstone, MD (Yale University) for helpful discussions. This work was funded by the National Institutes of Health (7U01AG084765-02). MS is additionally funded by the National Institutes of Health (2T32AG000029-49).

## AUTHOR CONTRIBUTIONS

Conceptualization: JO, AYV, MMS, AKD, SJ; Data Curation: MMS, AYV, AKD; Formal Analysis: AYV, MMS, AKD; Funding Acquisition: JO; Investigation: AYV, MMS, AKD; Project Administration: AYV, MMS, JO; Resources: JO, JS; Supervision: JO, SJ; Visualization and Writing: AYV, MMS; Writing - Review and Editing: MMS, AYV, AKD, JO, SJ.

## SUPPLEMENTAL TABLES AND LEGENDS

**Figure S1:**
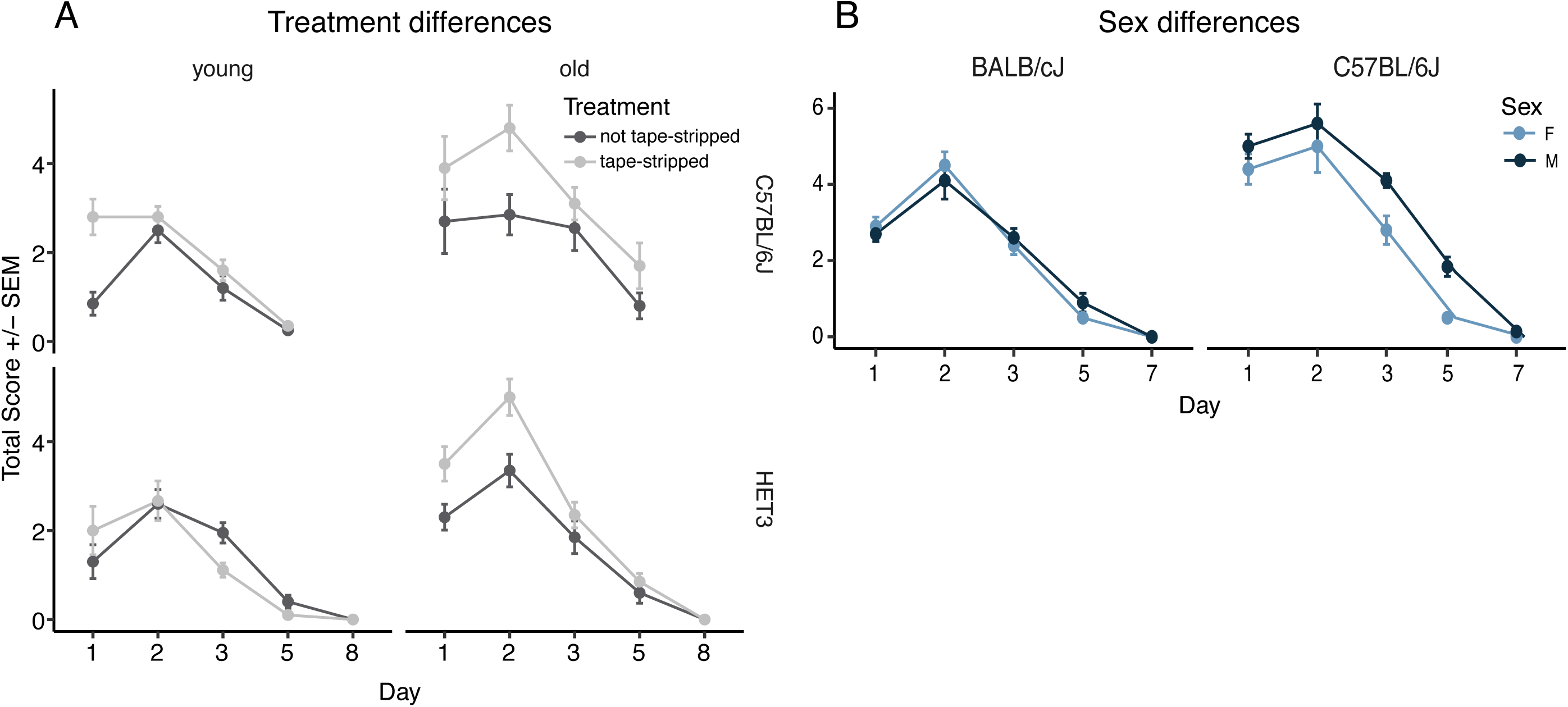
Additional parameters for model establishment. (**A**) Total lesion score with and without tape stripping assessed over time in young and old mice of inbred (C57BL/6J) versus outbred (HET3) strains (n(F)=10) showed modest strain or treatment-associated difference between healing trajectories. (**B**) Comparison of males versus females (BALB/cJ, n(M) = 5, n(F) = 5 (1F died before day 5) and C57BL/6J n(F) = 5, n(M) = 5, all tape-stripped) showed a concordant rate of lesion resolution with more severe lesions for males in C57BL/6J at 2 time points (day3 and 5). Tape-stripping experimental design is shown in Figure 1A. *signifies statistical significance with p<0.05; F: females, M: males

**Figure S2:**
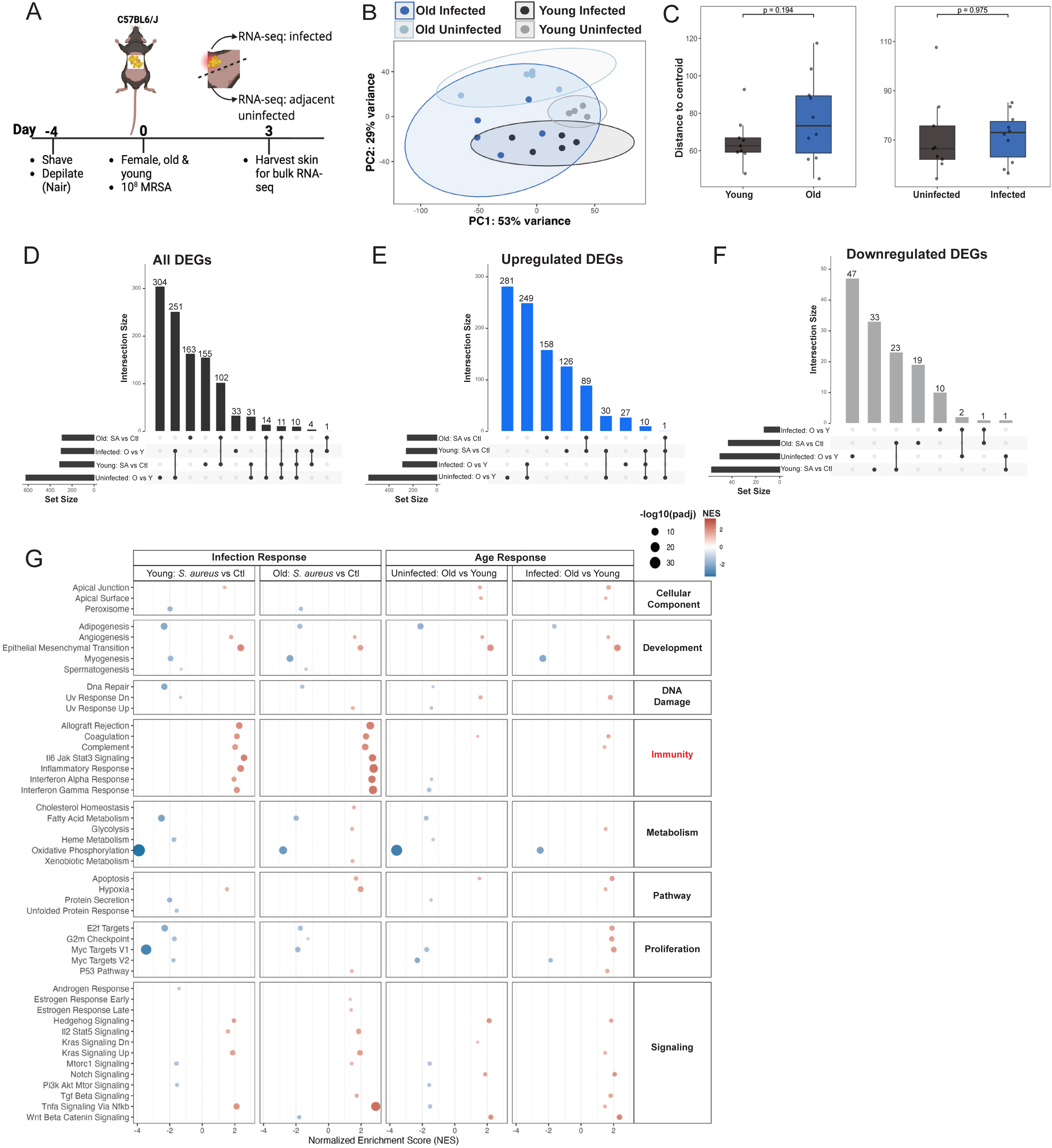
RNA sequencing reveals epidermal response to *S. aureus* infection in old and young animals. **(A)** In this first cohort, young (8 weeks) and old (80 weeks) female (n=5/age) C57BL/6J mice were epicutaneously exposed to *S. aureus* (1×10^8^ CFU) for 3 days, and a 2cm^2^ infected or adjacent uninfected skin biopsy was harvested. (**B**) Principal component analysis of variance stabilized data for all comparison groups showed age (PERMANOVA R^2^ = 0.21139, F = 6.0981, p=0.001) and infection (PERMANOVA R^2^ = 0.20173, F = 5.8196, p = 0.001) as independent drivers (age:infection PERMANOVA R^2^ = 0.02604, F = 0.7381, p = 0.615) of transcriptional response. (**C**) Centroid distance calculation showed higher but not significantly different variance in old animals compared to young (ns, Tukey test). (**D)** Upset plot of all differentially expressed genes (|Log2FC ≥ 2| and padj. ≤ 0.05) (**E**) and subset by upregulated or (**F**) downregulated gene sets showed significant overlap in DEGs between young and old infected skin as well as many unique DEGs per comparison. (**G**) Gene Set Enrichment analysis of Hallmark Genes (padj < 0.05) showed similar enrichment in infection related terms in both young and aged infected animals. Log2FC: Log2 fold change.

**Figure S3.**
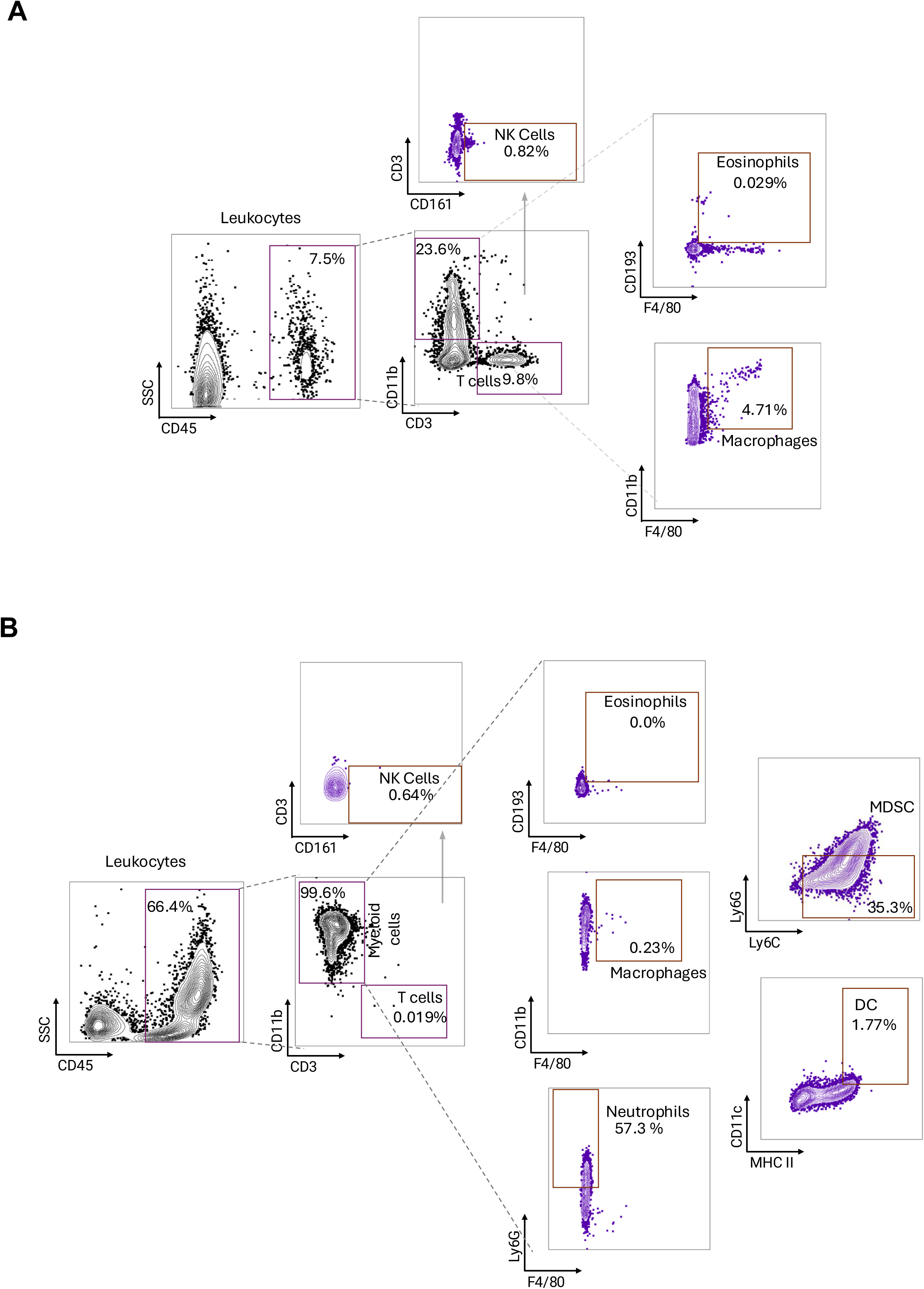
Flow cytometry gating strategy for immunophenotyping peripheral blood and microneedle patch samples. Representative sequential gating strategy used to identify live CD45⁺ immune cells and downstream myeloid and lymphoid populations in (**A**) peripheral blood samples and (**B**) cells recovered using microneedle patches. Immune populations included macrophages, neutrophils, dendritic cells, eosinophils, myeloid-derived suppressor cells (MDSCs), T cells and natural killer (NK) cells.

**Figure S4:**
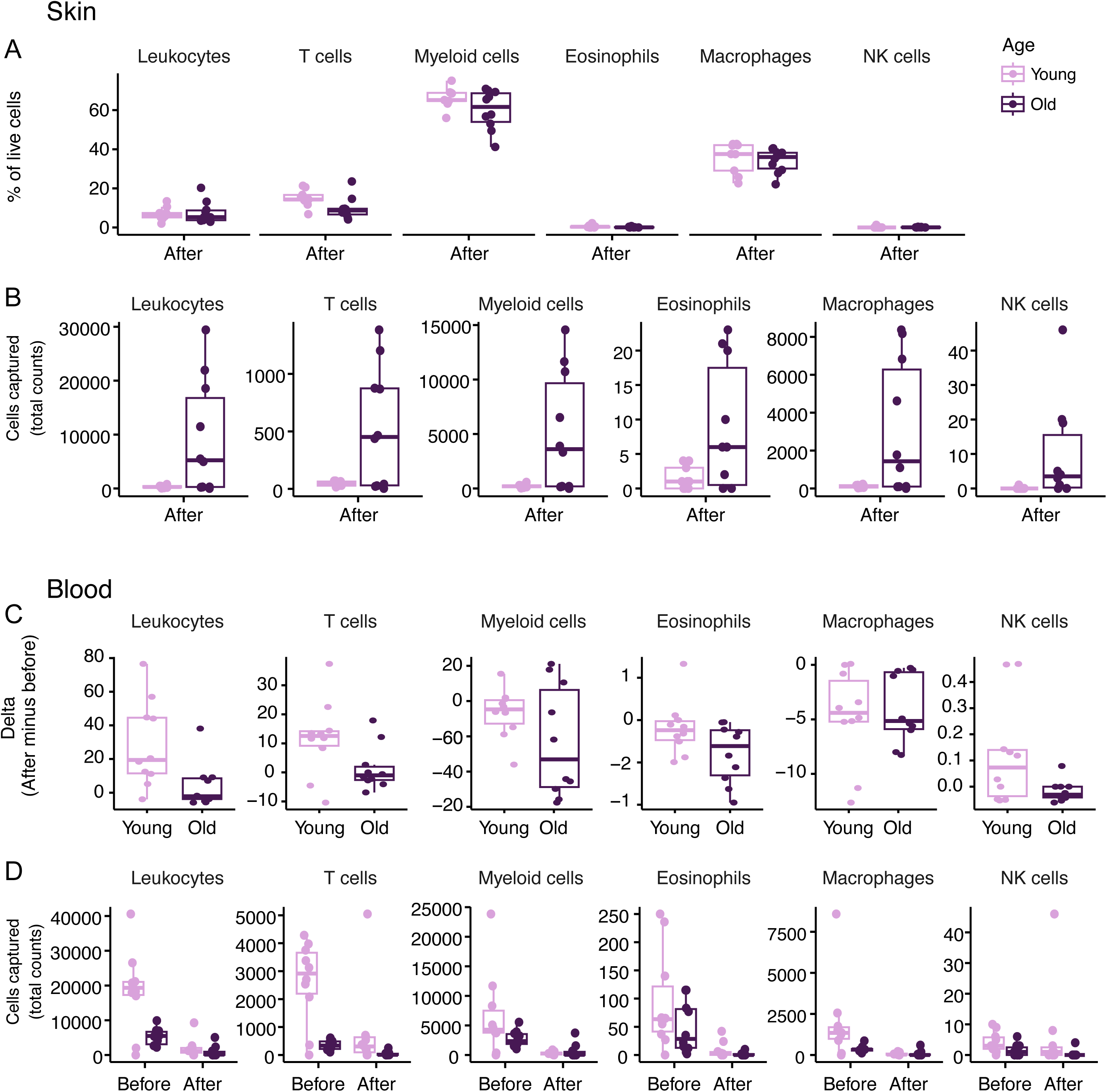
Cell counts detected in blood and skin biopsies. **(A)** Percent of live immune cells captured from infected skin biopsy tissue by flow cytometry showed no age associated differences in each immune subset. **(B)** total cell counts captured from infected skin biopsy tissue by flow cytometry at the experimental endpoint showed more total immune cells captured in aged tissues compared to young. **(C)** Change in immune cell counts in blood detected after infection compared to before infection (calculated as n(after) – n(before) showed no age associated systemic changes in each cell type. **(D)** Total detected cell counts detected by flow cytometry for each time point in retro-orbitally collected blood showed no age-associated difference in systemic cell counts. The input for cell counts from blood draws and skin biopsies could not be standardized; hence, no statistical evaluation was performed.

**Figure S5:**
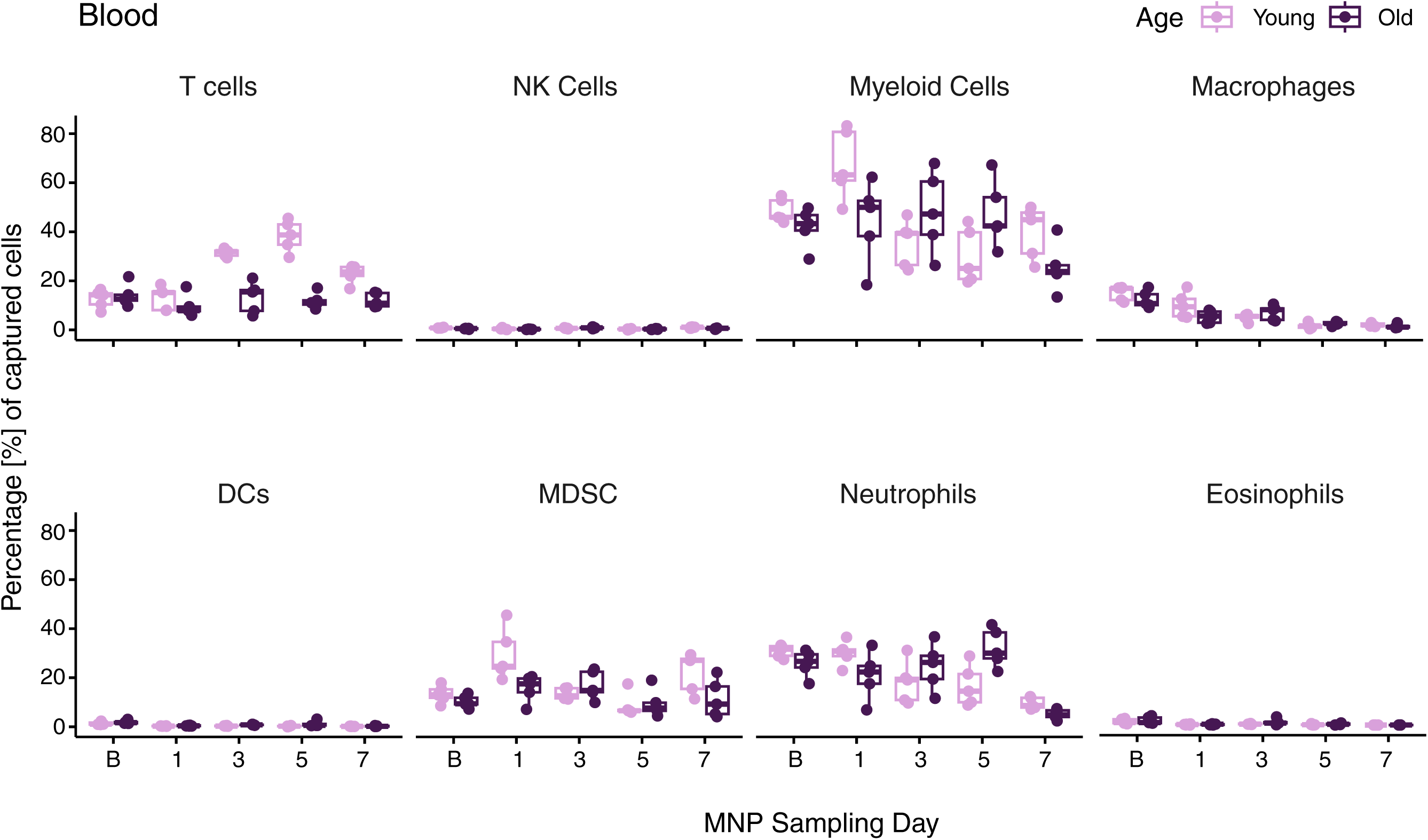
Immune cells captured from blood over time. **(A)** The percentage of each respective immune cell of the total captured cells is shown. Cells were identified by flow cytometry of retro-orbitally collected blood. Trajectory analysis revealed no significant differences in the change of each cell type over time between ages.

**Figure S6:**
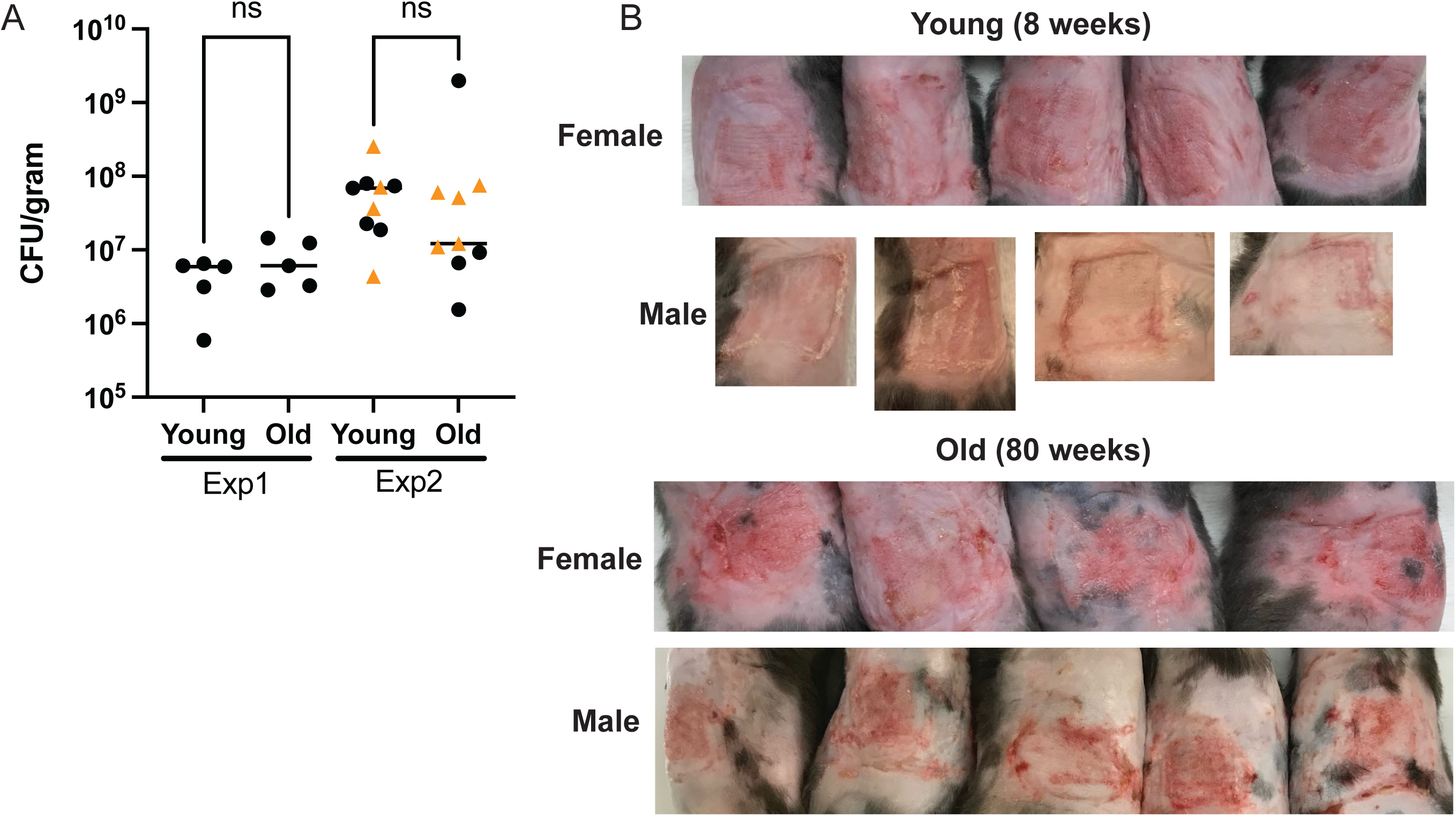
CFU counts and example images of infected skin. (**A**) CFU counts from infected tissue normalized to total grams recovered. Black circles are female animals; orange triangles are male animals. A non-parametric Kruskal-Wallis test followed by Dunn’s Multiple Comparisons Test was used to determine significant differences between groups. Experiment 1 refers to the tape stripping experiment from which skin was collected for RNA-seq where only non-tape stripped animals were included (Figure S2). Experiment 2 refers to the microneedle patch/RNA-seq paired experiment (Figures 2 and 4). No significant differences between ages or sexes were observed. (**B**) Representative images of lesions after 3-day infection and overnight application of microneedle patches. Tissue was then excised and processed for CFU quantification, RNA-seq, and Flow cytometry (Figures 2 and 4). Gross lesion severity was largely recapitulated across groups.

**Figure S7:**
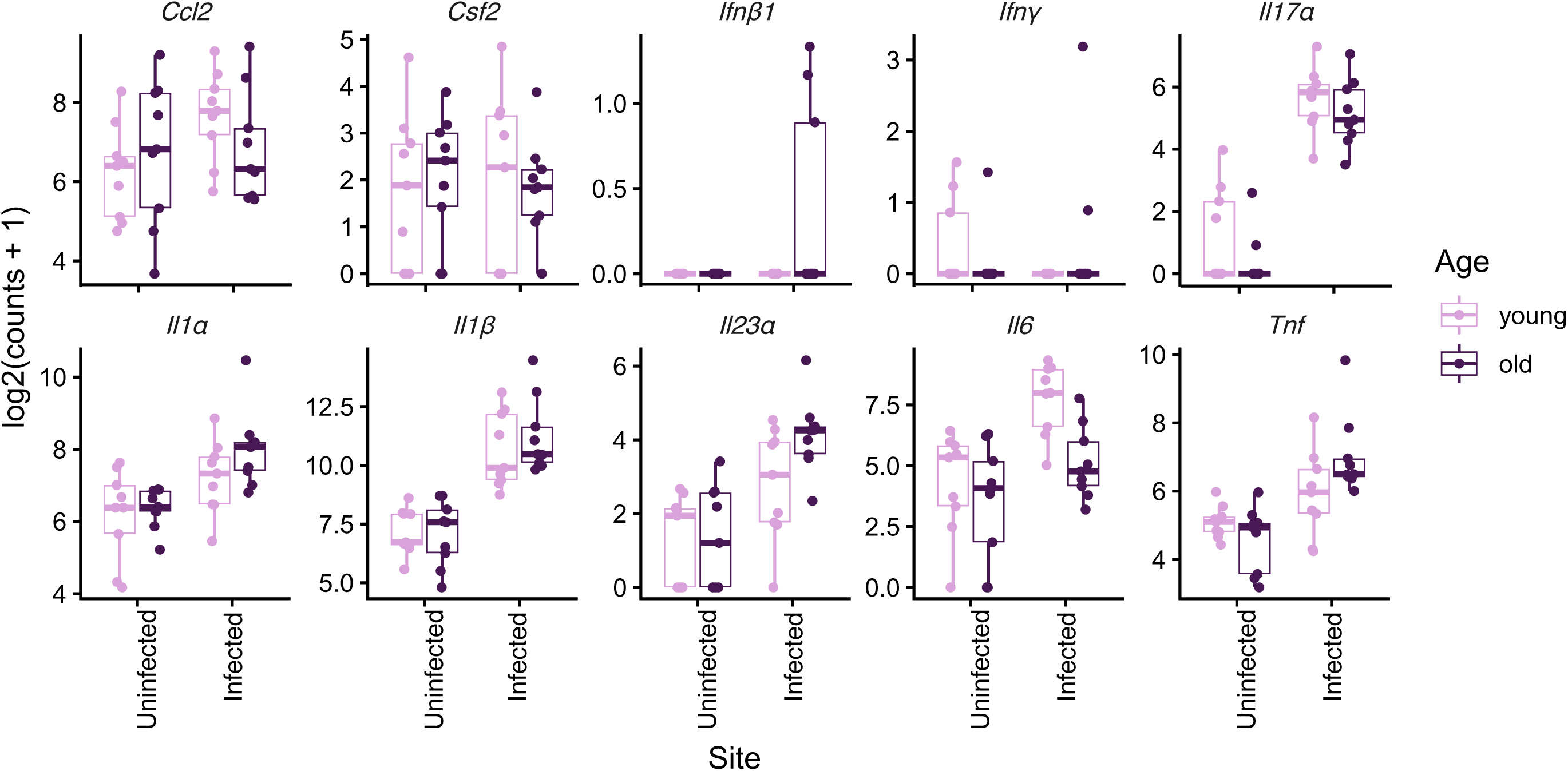
Transcript abundance of genes included in the cytokine panel. Deseq2 normalized gene counts for each transcript of genes included in the cytokine bead-based array. All counts were standardized as log2(count) + 1 to account for data points containing zero values. No significant differences between ages detected by Wilcox test for uninfected or infected tissue for any gene.

**Figure S8:**
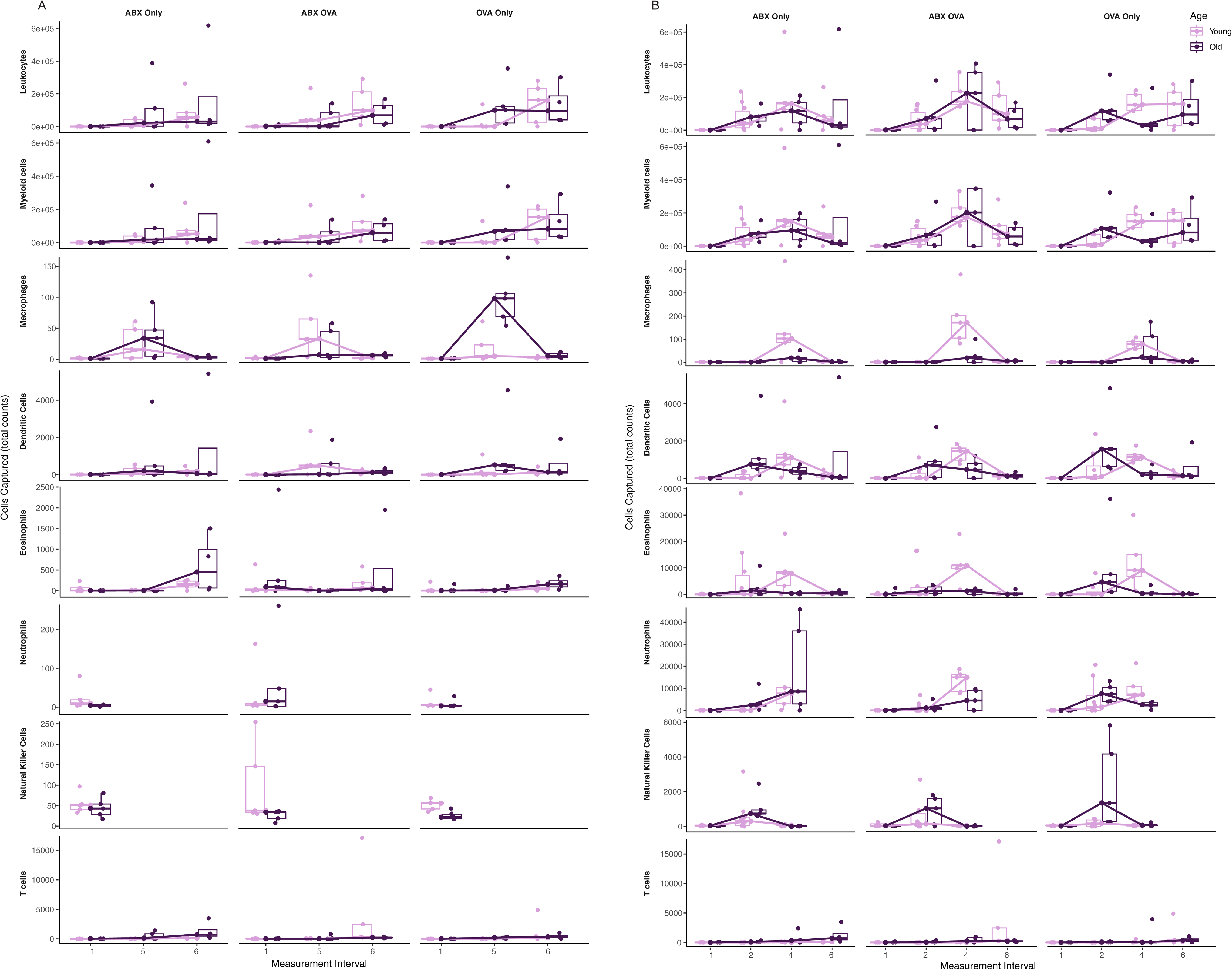
MNPs capture longitudinal immune response to microbiome depletion and OVA sensitization. Flow cytometry of MNP captured cells from each time point was plotted with either (**A**) an emphasis on cell counts pre-OVA treatment and end point or (**B**) an emphasis on post-OVA cell counts and endpoint. Captured cell data across groups showed time and treatment-dependent differences in immune cell capture in young versus old animals, all resolving by the final measurement interval. Measurement interval 1: Baseline; 2: After 1^st^ OVA; 3: Before 2^nd^ OVA (removed, technical issues); 4: After 2^rd^ OVA; 5: Before 3^rd^ OVA; 6: After 3^rd^ OVA (endpoint).

**Figure S9:**
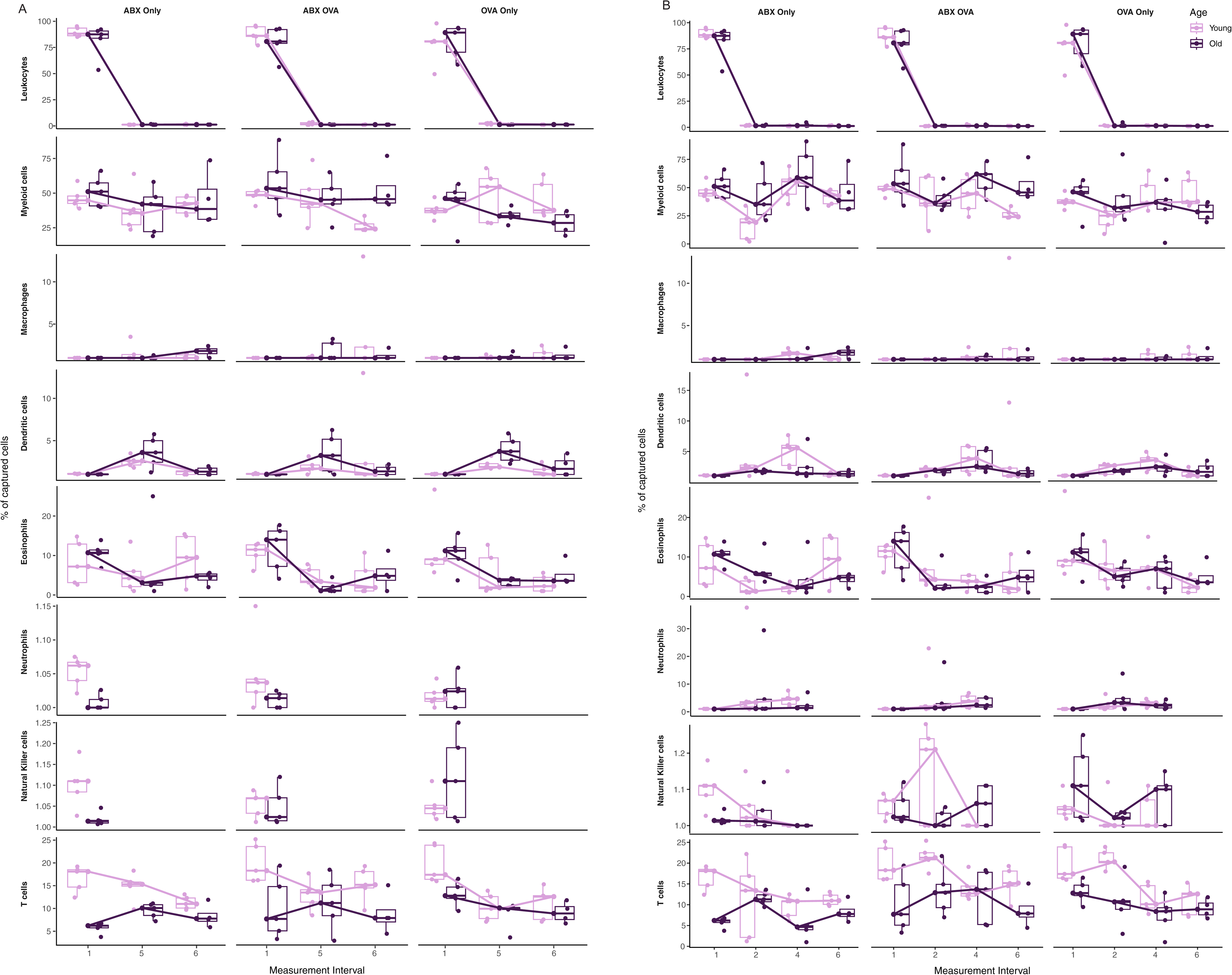
Cell percentages detected in blood in response to microbiome depletion and OVA sensitization. Flow cytometry of retro-orbitally collected blood from each time point was plotted with either (**A**) an emphasis on cell percentages in blood pre-OVA treatment and end point or (**B**) an emphasis on post-OVA cell percentages in blood and endpoint. Captured cell data across groups showed that systemic cell dynamics were decoupled from local dynamics captured by MNPs. Measurement interval 1: Baseline; 2: After 1^st^ OVA; 3: Before 2^nd^ OVA (removed, technical issues); 4: After 2^rd^ OVA; 5: Before 3^rd^ OVA; 6: After 3^rd^ OVA (endpoint).

**Figure S10:**
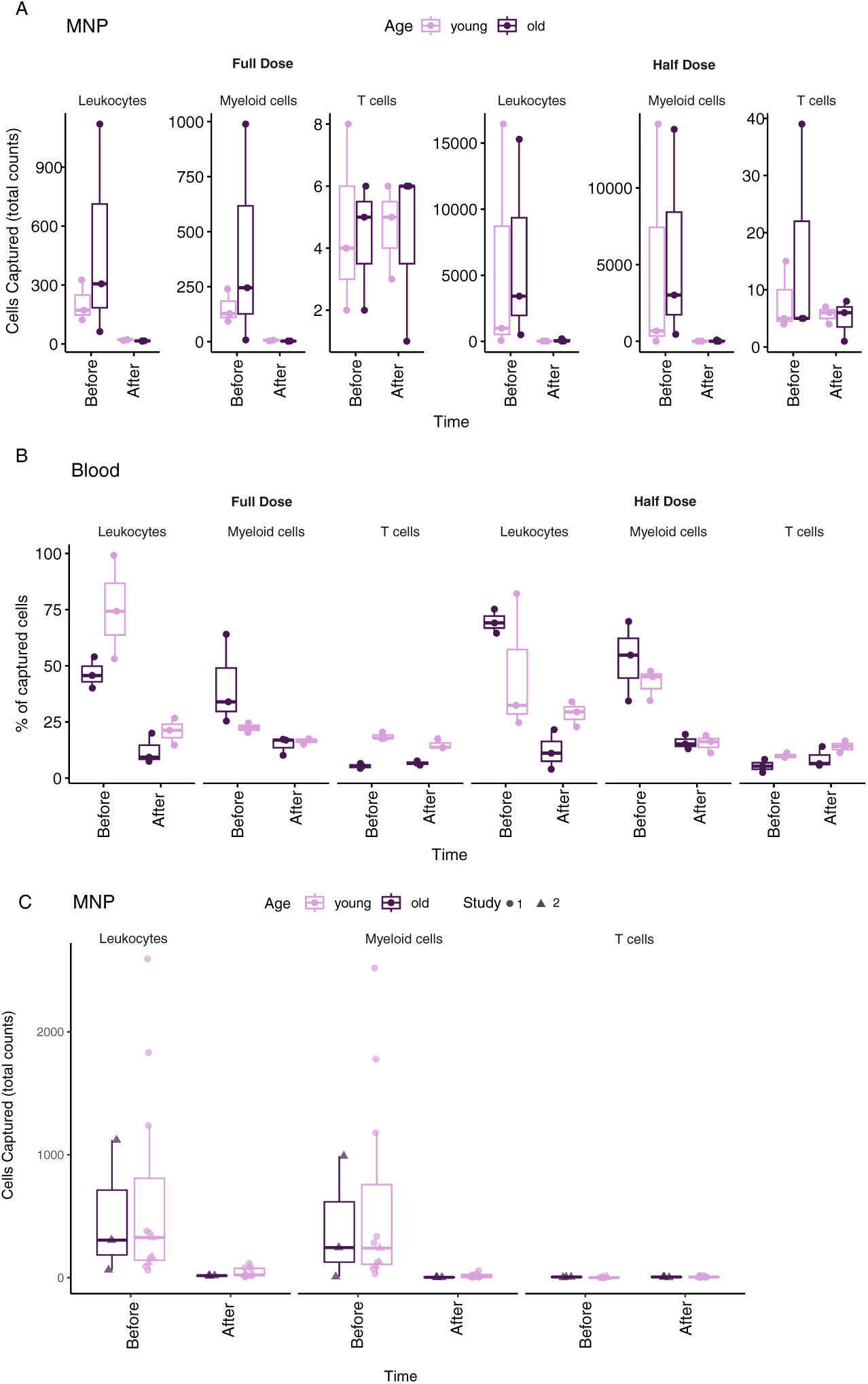
Select immune cells captured post systemic antibiotic treatment. **(A)** Flow cytometry of select immune cells captured with MNPs after systemic antibiotic treatment (full or half dose) in young and old mice showed antibiotic associated depletion of select immune cells in young and old animals. (**B**) Flow cytometry of select immune cells captured in reto-orbitally collected blood before and after systemic antibiotic treatment showed similar depletion of select immune cells systemically compared to MNPs. (**C**) Merge of MNP capture data for select immune cells following systemic antibiotic treatment in 2 cohorts of animals (denoted Study 1 and Study 2) showed reproducible effects between individual experiments.

## SUPPLEMENTAL TABLES AND LEGENDS

**Supplemental Table 1:** Median cytokine concentrations for young vs old mice in single time point experiment (related to Figure 2C). To calculate log2FC a pseudocount of 1 was added to normalize 0 (pg/mL pre infection) values. LOD: limit of detection.

| Cytokine | Young post infection concentration (median pg/mL) | Young log<sub>2</sub>FC (post vs pre infection) | Old post infection concentration (median pg/mL) | Old log<sub>2</sub>FC (post vs pre infection) |
| --- | --- | --- | --- | --- |
| GM-CSF | 51.28 | 5.7 | 30.97 | 5.0 |
| IL17-a | 81.18 | 6.4 | 84.71 | 6.4 |
| IL-1a | >16000 (LOD) | 2.2 | 13708.14 | 3.6 |
| IL-1B | 315.34 | 8.3 | 228.85 | 7.8 |
| TNF-a | 124.35 | 7.0 | 101.31 | 6.7 |

**Supplemental Table 2:** Significant age associated differences in T cell numbers captured by MNPs pre and post infection for single time point experiment (related to Figure 2D).

| Cell Type | Young pre infection (cell count) | Old pre infection (cell count) | Pre infection difference in medians (Wilcox test) | Young post infection (cell count) | Old post infection (cell count) | Post infection difference in medians (unpaired Wilcox) |
| --- | --- | --- | --- | --- | --- | --- |
| MNP T-cells | Median = 4<br>Mean = 150 | Median = 104.5<br>Mean = 109 | P=0.014 | Median = 10994.0<br>Mean = 13757.8 | Median = 256.0<br>Mean = 680.3 | P=0.00052 |

**Supplemental Table 3:** Age associated differences in T cell and leukocyte percentages in blood pre and post infection for single time point experiment (related to Figure 2E).

| Cell Type | Young pre infection (%) | Old pre infection (%) | Young post infection (%) | Old post infection (%) |
| --- | --- | --- | --- | --- |
| T cells (blood) | Median = 13.71<br>Mean = 13.48 | Median = 6.70<br>Mean = 6.75 | Median = 25.6<br>Mean = 25.3 | Median = 5.01<br>Mean = 8.21 |
| Leukocytes (blood) | ND | Median = 23.78<br>Mean = 33.57 | ND | Median = 1.93<br>Mean = 8.01 |

## REFERENCES

1. Choi, E. H. Aging of the skin barrier. Clin. Dermatol. 37, 336–345 (2019).

2. Farage, M. A., Miller, K. W., Elsner, P. & Maibach, H. I. Characteristics of the Aging Skin. Adv. Wound Care 2, 5–10 (2013).

3. Laube, S. Skin infections and ageing. Ageing Res. Rev. 3, 69–89 (2004).

4. Pilkington, S. M., Bulfone-Paus, S., Griffiths, C. E. M. & Watson, R. E. B. Inflammaging and the Skin. J. Invest. Dermatol. 141, 1087–1095 (2021).

5. Jafferany, M., Huynh, T. V., Silverman, M. A. & Zaidi, Z. Geriatric dermatoses: a clinical review of skin diseases in an aging population. Int. J. Dermatol. 51, 509–522 (2012).

6. Falcone, M. & Tiseo, G. Skin and soft tissue infections in the elderly. Curr. Opin. Infect. Dis. 36, 102–108 (2023).

7. Linz, M. S., Mattappallil, A., Finkel, D. & Parker, D. Clinical Impact of Staphylococcus aureus Skin and Soft Tissue Infections. Antibiot. Basel Switz. 12, 557 (2023).

8. Ray, G. T., Suaya, J. A. & Baxter, R. Incidence, microbiology, and patient characteristics of skin and soft-tissue infections in a U.S. population: a retrospective population-based study. BMC Infect. Dis. 13, 252 (2013).

9. Kang, C.-I., Song, J.-H., Ko, K. S., Chung, D. R. & Peck, K. R. Clinical features and outcome of Staphylococcus aureus infection in elderly versus younger adult patients. Int. J. Infect. Dis. 15, e58–e62 (2011).

10. Kennedy, A. D. et al. Epidemic community-associated methicillin-resistant *Staphylococcus aureus* : Recent clonal expansion and diversification. Proc. Natl. Acad. Sci. 105, 1327–1332 (2008).

11. Youn, C., Archer, N. K. & Miller, L. S. Research Techniques Made Simple: Mouse Bacterial Skin Infection Models for Immunity Research. J. Invest. Dermatol. 140, 1488–1497.e1 (2020).

12. Patrick, G. J. et al. Epicutaneous Staphylococcus aureus induces IL-36 to enhance IgE production and ensuing allergic disease. J. Clin. Invest. 131, e143334 (2021).

13. Steinbach, S. et al. Temporal dynamics of intradermal cytokine response to tuberculin in Mycobacterium bovis BCG-vaccinated cattle using sampling microneedles. Sci. Rep. 11, 7074 (2021).

14. Samant, P. P. et al. Sampling interstitial fluid from human skin using a microneedle patch. Sci. Transl. Med. 12, eaaw0285 (2020).

15. Churchill, G. A., Gatti, D. M., Munger, S. C. & Svenson, K. L. The Diversity Outbred Mouse Population. Mamm. Genome Off. J. Int. Mamm. Genome Soc. 23, 713–718 (2012).

16. Bitschar, K. et al. *Staphylococcus aureus* Skin Colonization Is Enhanced by the Interaction of Neutrophil Extracellular Traps with Keratinocytes. J. Invest. Dermatol. 140, 1054–1065.e4 (2020).

17. Wanke, I. et al. Staphylococcus aureus skin colonization is promoted by barrier disruption and leads to local inflammation. https://doi.org/10.1111/exd.12083 doi:10.1111/exd.12083.

18. Kugelberg, E. et al. Establishment of a Superficial Skin Infection Model in Mice by Using Staphylococcus aureus and Streptococcus pyogenes. Antimicrob. Agents Chemother. 49, 3435–3441 (2005).

19. Quiros-Roldan, E., Sottini, A., Natali, P. G. & Imberti, L. The Impact of Immune System Aging on Infectious Diseases. Microorganisms 12, 775 (2024).

20. Jalili, S. et al. Leveraging tissue-resident memory T cells for non-invasive immune monitoring via microneedle skin patches. *Nat*. Biomed. Eng. 1–16 (2026) doi:10.1038/s41551-026-01617-7.

21. Wang, G. et al. Repeated epicutaneous exposures to ovalbumin progressively induce atopic dermatitis-like skin lesions in mice. Clin. Exp. Allergy 37, 151–161 (2007).

22. Jin, H., He, R., Oyoshi, M. & Geha, R. Animal models of atopic dermatitis. J. Invest. Dermatol. 129, 31–40 (2009).

23. Hasanpour, A. H. et al. The global prevalence of methicillin-resistant Staphylococcus aureus colonization in residents of elderly care centers: a systematic review and meta-analysis. Antimicrob. Resist. Infect. Control 12, 4 (2023).

24. Larson, P. J. et al. Associations of the skin, oral and gut microbiome with aging, frailty and infection risk reservoirs in older adults. *Nat*. Aging 2, 941–955 (2022).

25. Mendes, A. I., Piexoto, M. J., Marques, A. P., Pedrosa, J. & Fraga, A. G. An optimized mouse model of Staphylococcus aureus infected diabetic ulcers. BMC Res. Notes 15, 293 (2022).

26. Kane, A. E. et al. Animal models of frailty: current applications in clinical research. Clin. Interv. Aging 11, 1519–1529 (2016).

27. Kimmel, J. C. et al. Murine single-cell RNA-seq reveals cell-identity- and tissue-specific trajectories of aging. Genome Res. 29, 2088–2103 (2019).

28. Zonnefeld, A. G. et al. Characterization of age-associated gene expression changes in mouse sweat glands. Aging 16, 6717–6730 (2024).

29. Wen, S. et al. Aged and young mice differentially respond to tape-stripping in epidermal gene expression. Exp. Dermatol. 31, 312–319 (2022).

30. Kline, S. N., et al. *Staphylococcus aureus* proteases trigger eosinophil-mediated skin inflammation. Proc. Natl. Acad. Sci. 121, e2309243121 (2024).

31. Liu, H. et al. Staphylococcus aureus epicutaneous exposure drives skin inflammation via IL- 36-mediated T cell responses. Cell Host Microbe 22, 653–666.e5 (2017).

32. Nakagawa, S. et al. Staphylococcus aureus Virulent PSMα Peptides Induce Keratinocyte Alarmin Release to Orchestrate IL-17-Dependent Skin Inflammation. Cell Host Microbe 22, 667–677.e5 (2017).

33. Tseng, C. W. et al. Innate Immune Dysfunctions in Aged Mice Facilitate the Systemic Dissemination of Methicillin-Resistant S. aureus. PLoS ONE 7, e41454 (2012).

34. Nagarajan, A. et al. Collaborative Cross mice have diverse phenotypic responses to infection with Methicillin-resistant Staphylococcus aureus USA300. PLOS Genet. 20, e1011229 (2024).

35. Castleman, M. J. et al. Innate Sex Bias of Staphylococcus aureus Skin Infection Is Driven by α-Hemolysin. J. Immunol. 200, 657–668 (2018).

36. Pomorska-Wesołowska, M. et al. Longevity and gender as the risk factors of methicillin- resistant Staphylococcus aureus infections in southern Poland. BMC Geriatr. 17, 51 (2017).

37. John, M., Chinnappan, M., Sturges, C. & Harris-Tryon, T. Skin androgens regulate Staphylococcus aureus pathogenicity via quorum sensing. 11, 701–717 (2026).

38. Wilkening, R. V., Langouët-Astrié, C., Severn, M. M., Federle, M. J. & Horswill, A. R. Identifying genetic determinants of Streptococcus pyogenes-host interactions in a murine intact skin infection model. Cell Rep. 42, 113332 (2023).

39. Kovacs, E. J. et al. Aging and innate immunity in the mouse: impact of intrinsic and extrinsic factors. Trends Immunol. 30, 319–324 (2009).

40. Brubaker, A. L., Rendon, J. L., Ramirez, L., Choudhry, M. A. & Kovacs, E. J. Reduced Neutrophil Chemotaxis and Infiltration Contributes to Delayed Resolution of Cutaneous Wound Infection with Advanced Age. J. Immunol. 190, 1746–1757 (2013).

41. Faraji Rad, Z., Prewett, P. D. & Davies, G. J. An overview of microneedle applications, materials, and fabrication methods. Beilstein J. Nanotechnol. 12, 1034–1046 (2021).

42. Sakamoto, K. & Nagao, K. Mouse Models for Atopic Dermatitis. Curr. Protoc. 3, e709 (2023).

43. Wickham, H. et al. Welcome to the Tidyverse. J. Open Source Softw. 4, 1686 (2019).

44. Wickham, H. Ggplot2. (Springer International Publishing, Cham, 2016). doi:10.1007/978-3-319-24277-4.

45. Kassambara, A. ggpubr: ‘ggplot2’ Based Publication Ready Plots. R package version 0.6.3. https://rpkgs.datanovia.com/ggpubr/ https://rpkgs.datanovia.com/ggpubr/authors.html#citation (2026).

46. Wickham, H. Reshaping Data with the reshape Package. J. Stat. Softw. 21, 1–20 (2007).

47. Bates, D., Mächler, M., Bolker, B. & Walker, S. Fitting Linear Mixed-Effects Models Using lme4. J. Stat. Softw. 67, 1–48 (2015).

48. McGillycuddy, M., Popovic, G., Bolker, B. M. & Warton, D. I. Parsimoniously Fitting Large Multivariate Random Effects in glmmTMB. J. Stat. Softw. 112, 1–19 (2025).

49. Lenth, R. V. et al. emmeans: Estimated Marginal Means, aka Least-Squares Means. (2026).

50. Uberoi, A. et al. Commensal microbiota regulates skin barrier function and repair via signaling through the aryl hydrocarbon receptor. Cell Host Microbe 29, 1235–1248.e8 (2021).

51. Bolger, A. M., Lohse, M. & Usadel, B. Trimmomatic: a flexible trimmer for Illumina sequence data. Bioinformatics 30, 2114–2120 (2014).

52. Dobin, A. et al. STAR: ultrafast universal RNA-seq aligner. Bioinformatics 29, 15–21 (2013).

53. Liao, Y., Smyth, G. K. & Shi, W. featureCounts: an efficient general purpose program for assigning sequence reads to genomic features. Bioinformatics 30, 923–930 (2014).

54. Risso, D., Ngai, J., Speed, T. P. & Dudoit, S. Normalization of RNA-seq data using factor analysis of control genes or samples. Nat. Biotechnol. 32, 896–902 (2014).

55. Love, M. I., Huber, W. & Anders, S. Moderated estimation of fold change and dispersion for RNA-seq data with DESeq2. Genome Biol. 15, 550 (2014).

56. vegan: an R package for community ecologists. https://vegandevs.github.io/vegan/.

57. pheatmap: Pretty Heatmaps. https://raivokolde.r-universe.dev/pheatmap.

58. Conway, J. R., Lex, A. & Gehlenborg, N. UpSetR: an R package for the visualization of intersecting sets and their properties. Bioinformatics 33, 2938–2940 (2017).

59. Yu, G., Wang, L.-G., Han, Y. & He, Q.-Y. clusterProfiler: an R Package for Comparing Biological Themes Among Gene Clusters. OMICS J. Integr. Biol. 16, 284–287 (2012).

60. Wu, T. et al. clusterProfiler 4.0: A universal enrichment tool for interpreting omics data. The Innovation 2, 100141 (2021).

